# Characterizing the Assembly and Functional Properties of Gene-Length Mixed DNA Monolayers on Electrodes for Cell-Free Expression

**DOI:** 10.64898/2026.08.05.742923

**Authors:** Rory Majule, Kyvalya Reddy, Samrutha Babu, Oren Fox, Jeff Nivala, Chris Takahashi

## Abstract

Gold electrodes are attractive substrates for bioelectronic and cell-free synthetic biology platforms because they are conductive, chemically stable, biocompatible, and readily functionalized through thiol-gold chemistry. Here, gene-length DNA monolayers assembled on planar gold electrodes as reusable templates for cell-free protein expression are investigated.

Using thiol-modified sfGFP genes, DNA surface density is shown to be tunable by changing the DNA concentration during incubation, with the immobilized genes able to support cell-free sfGFP expression directly from the electrode surface. Further, the effects of applied voltage, storage, repeated reactions, reducing agents, and protein fouling on monolayer stability and expression output are examined. While some conditions lead to loss of reusable expression activity, dense chemisorbed monolayers can retain partial function under neutral, non-reducing conditions and are relatively robust to protein exposure. In contrast, low-density physisorbed monolayers show a stronger relationship between DNA loss and expression output. Finally, when using gold-mediated fluorescence quenching to monitor changes in DNA conformation, surface-bound DNA demonstrates electrophoretic addressability. Together, these results establish DNA- functionalized planar electrodes as a promising foundation for modular, addressable cell-free expression platforms.

## 1 Introduction

Cell-free gene expression systems can be used in the development of hybrid functional materials, to design synthetic biological and biocomputational systems, and to reduce biological complexity when investigating genetic pathways found in nature.^[1]^ While cell free expression (CFE) reactions can be carried out in tubes or in microwell plates, any components of the reaction that would be reused in a cellular environment, such as DNA, must instead be replenished every time reactions are repeated or iterated upon. The level of control one has over the timing and dynamics of the reaction is also largely limited to the relative concentrations of each added component and to the tuning of environmental factors such as incubation temperature. Finally, while expression dynamics within a cell are governed in part by the spatial separation of genetic material from the bulk reaction occurring in the cytoplasm, as well as by diffusion dynamics between multiple gene elements, these spatial dynamics cannot be easily replicated in a cell-free context without the use of some material platform. To these ends, gel-based platforms^[2]^ as well as platforms using silicon dioxide-bound DNA brushes^[3–6]^ have sought to remedy the issues of DNA reuse as well as to impart some degree of control over reaction dynamics through the geometric separation of genetic components. While such platforms certainly allow for more nuanced control of CFE reactions when compared to the more traditional test-tube setup, the desired diffusion dynamics must be “hard-coded” into the system. In other words, these systems lack addressability once the platform has been fabricated and the reaction initiated. This contrasts with *in vivo* transcription and translation. A variety of reactions and genetic circuits can be expressed within the cell at any given time, and regulation of transcription and translation can occur in response to external factors while gene expression is taking place. Living cells also harness spatially driven regulation involving symmetry breaking and both intra- and intercell diffusion, all of which would be difficult to implement in bulk solutions. Microfluidic chips with switching valves and addressable reaction chambers^[7]^ have enabled both steady-state and dynamic control of protein synthesis reactions;^[8]^ however, their platform complexity scales with the number of reactions being implemented.

Implementation on an electronically addressable surface could enable a unique range of functionalities not readily accessible in conventional cell-free expression platforms. For example, a DNA-modified electrode platform could integrate CFE and aptameric ligand sensing,^[9,10]^ with aptamers either embedded within a gene brush to generate real-time, electrochemically measured expression curves, or situated adjacent to a gene brush on the same chip to take advantage of passive diffusion. Beyond electrochemical sensing, performing cell-free transcription-translation (TX-TL) reactions on an electrode platform could also dynamic control of expression through electrochemically driven hybridization, strand melting,^[11]^ or local pH control.^[12,13]^

While a number of studies have been published demonstrating CFE using gene fragments immobilized on or in electronically inert interfaces^[2–6]^ or demonstrating other enzymatic reactions such as polymerase chain reactions (PCR) using DNA-modified electrodes as a platform,^[14]^ to our knowledge none have demonstrated the feasibility of carrying out CFE reactions using gene fragments chemically bound to a gold surface. To investigate this, we developed a system to chemically attach gene-encoding dsDNA to a gold electrode surface using conventional gold-thiol chemistry. We found that these biochips retained functionality for weeks when stored at 4 ℃ and that these self-assembled monolayers (SAMs) had a predictable shelf-life that was dependent on packing density, storage time, and number of CFE cycles. We also examined the way biofouling layers interact with the monolayer, with results suggesting that low-density physisorbed monolayers are more easily disrupted by biofouling than higher density, chemisorbed monolayers. Lastly, we confirmed that conformational manipulation of the surface-bound strands can be controlled electrochemically, and that this conformational control is possible even when the surface-bound dsDNA is significantly longer than the persistence length.

## 2 Materials and Methods

### DNA Fragments and Oligos

Double-stranded DNA (dsDNA) fragments 968 base pairs (bp) in length containing the super-folder green fluorescent protein (sfGFP) gene with a T7 promoter (**Table S1**) were purchased from Integrated DNA Technologies (IDT). This DNA fragment was used as a template in PCR to add modifications, such as a 5′ thiol C6 S-S and/or a 5′ Cy3 fluorophore using modified primers purchased from IDT. When the 5′ thiol group was attached to the forward primer, the thiol was stated to be on the “proximal” end, since it was located proximal to the promoter region, and the resulting modified gene fragment was referred to as “outward facing” in reference to the direction a T7 RNA polymerase (T7 RNAP) would travel along the strand relative to the electrode surface. Inversely, when the 5′ thiol group was attached to the reverse primer, the thiol was stated to be on the “distal” end, and the resulting modified gene fragment was referred to as “inward facing.” Modified and unmodified PCR products were purified using either the Monarch PCR & DNA Cleanup Kit (T1030, New England Biolabs) or DINOMAG Clean-Up & Size Selection SPRI Magnetic Beads (DN9004, LabScoop). All purified fragments and duplexes were eluted and stored in molecular grade water.

For microscopy experiments, a 72mer oligonucleotide (from Kaiser and Rant^[15]^) designed to minimize the occurrence of secondary structures: 5′-HS-(CH_2_)_6_-TAG TCG TGA GCA CAT GGA CCT GAT TAG TCG TAA GCT GAT ATG GCT GAT TAG TCG GAA GCA TCG AAC GCT GAT-Cy3-3′. Before immobilization on a gold electrode surface, a duplex was formed by hybridizing to to a reverse complement strand. Hybridization was performed by mixing the modified oligo and an excess of its unmodified reverse complement in annealing buffer (1 mM EDTA, 10 mM Tris, 50 mM NaCl) and slowly ramping down from 90 ℃ to 25 ℃. Excess, unannealed oligos were digested in Exonuclease I (M0293S, New England Biolabs) and then purified using the oligo cleanup protocol of the Monarch PCR & DNA Cleanup Kit.

### Gold Electrode Preparation

Gold coated microscope slides were either purchased from Substrata (SKU SSAU1000) or were fabricated at the Washington Nanofabrication Facility (WNF). In both cases, gold layers were deposited onto borosilicate glass with thickness of 1000 Å for Substrata slides and 500 Å for WNF-fabricated slides, with a 2-7 nm thick chromium adhesion layer using high vacuum electron beam evaporation. Macro-scale patterning was achieved on the 500 Å slides by applying laser-cut Kapton tape to the glass slides prior to evaporation. Gold layers were confirmed to be predominately of Au(111) crystal orientation using a Bruker D8 Discover x-ray diffractometer (**Figure S1**). Immediately prior to first use, gold slides were chemically cleaned via a 10 minute immersion in Nano-Strip^®^ (KMG Electronic Chemicals, Inc.) warmed to > 60 ℃. The slides were then rinsed three times in molecular grade (MilliQ) water, dried under argon gas, and then plasma cleaned using O_2_ plasma in a Harrick Plasma PDC-001-HP plasma cleaner.^[16]^ During O_2_ plasma cleaning, pressure was maintained between 600 and 800 mTorr for five minutes with the radio frequency power level set to approximately 45 W.

Alternatively, when specified, gold disk electrodes (CHI101, CH Instruments) were used as substrate surfaces. Prior to use, disk electrodes were polished first with 1.0, then 0.3, then 0.05 micron alumna slurry for four minutes each. Residual alumina powder was removed by placing electrodes in a secondary vessel suspended in a degassed Branson CPX2800H ultrasonic bath.

The secondary vessel was filled with 100% ethanol and processed for 10 minutes and then replaced with MilliQ water and processed for another 10 minutes. Immediately before monolayer immobilization, residual impurities were removed through electrochemical oxidation and reduction. A two segment scan from −1 to −1.8 V vs Ag/AgCl was performed in 0.5 M sodium hydroxide (NaOH) 300 times at a scan rate of 1 V s^−1^. A three segment scan from +0.4 to −0.5 to +1.75 V and back to +0.4 V vs Ag/AgCl was then performed in 0.5 M sulfuric acid (H_2_SO_4_) 200 times at a scan rate of 1 V s^−1^. Cleanness was checked by running a single cyclic voltammetry (CV) cycle in fresh 0.5 M H_2_SO_4_ with a potential range of -0.3 to 1.55 V versus Ag/AgCl, and a scan rate 0.1 V s^−1^.^[17]^

### Preparing DNA for Immobilization

Purified linear sfGFP gene fragments were concentrated in a Thermo Scientific Savant DNA120 SpeedVac Concentrator until the desired concentration was reached. The thiol groups were reduced by performing a one-to-one dilution of the aqueous DNA in a solution of 2 mM tris(2-carboxyethyl)phosphine (TCEP), 10 mM Tris-HCl, and 15 mM NaCl. The resulting solution was left in the dark for approximately 2 hours unless otherwise stated, after which the DNA solution was further diluted in four parts immobilization buffer. The immobilization buffer was a solution of 10 mM Tris-HCl (pH 7.5), 15 mM NaCl, and 50 mM MgCl_2_.

### Immobilizing DNA on Gold Electrodes

i-well microscope slide chambers (ProPlate^®^) were obtained from Grace Bio-Labs to subdivide the gold-coated microscope slides into multiple, leak-proof, reagent wells without the use of adhesive. The slide chambers consisted of a silicone gasket sheet, a plastic multi-well module, and clips to hold the components secularly in place on the surface of the microscope slide. Prior to use or reuse, the plastic module and silicone gasket were scrubbed in diluted Liquinox detergent, rinsed in deionized water, and then ultrasonically cleaned in both 100% ethanol and 100% isopropyl alcohol for 10 minutes each. Both the plastic and silicone components were then autoclaved to ensure removal of DNase contamination.

When preparing multiple monolayers on a single gold coated slide, a clean ProPlate chamber was secured to the slide and 10 or 30 µl of the DNA or control solution was aliquoted into the chamber and onto the cleaned gold surface. The volume used was dependent on the surface area of the slide chambers to ensure total surface coverage. For immobilization on disk electrodes, 5 µl of the DNA or control solution was pipetted onto the gold disk, which was then capped. Unless otherwise stated, the DNA or control solution was then frozen on the gold surface before being removed from the freezer and being allowed to sit at room temperature for 16-24 hours. After incubation, monolayer areas were rinsed several times in 10 mM Tris-HCl and then immersed in 1 mM 6-mercapto-1-hexanol (MCH) in immobilization buffer for 1 hour to remove physisorbed DNA molecules (Error! Reference source not found.). After MCH incubation, electrodes were again rinsed in both Tris-HCl and phosphate-buffered saline (PBS). Unless otherwise specified, monolayers were stored in 1X PBS at 4 ℃ until ready to use.

**Scheme 1.**
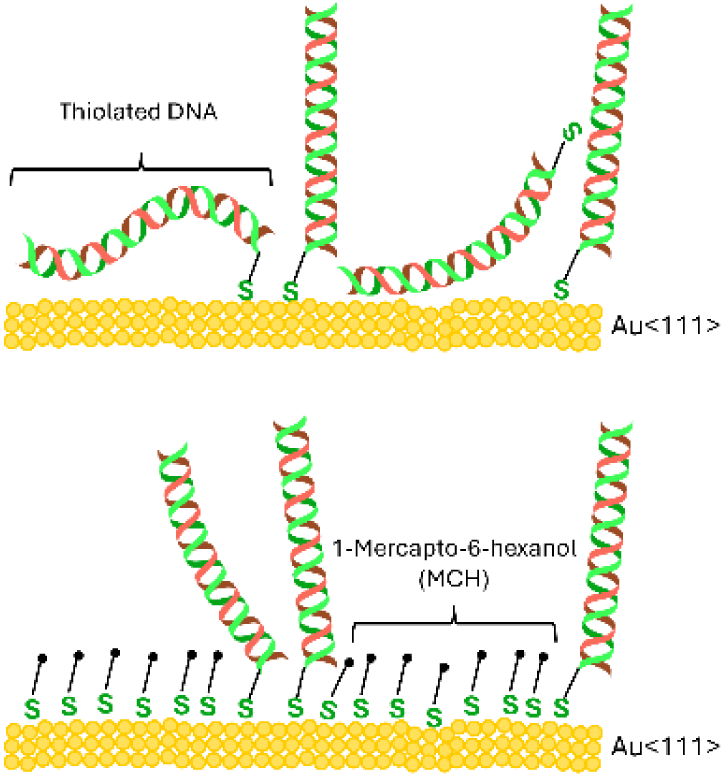
968 bp-long dsDNA with a 5′ C6 thiol-modification is immobilized on the gold electrode surface. MCH is incubated on the surface to backfill gaps and displace physisorbed DNA molecules. This procedure is also performed using unmodified dsDNA to evaluate the effectiveness of the MCH backfilling and surface washing steps.

### Chronocoulometric Quantification of Monolayer Packing Density

Chronocoulometry (CC) was employed to quantify the amount of DNA strands immobilized at gold surfaces as previously reported.^[17–19]^ Individual DNA/MCH-modified gold electrodes were connected to a WaveNow XV potentiostat (Pine Research) running with AfterMath software, and were immersed in 10 mM Tris-HCl (pH 7.4) along with a Ag/AgCl reference electrode (CHI111, CH Instruments, stored in 1 M KCl) and a Pt wire counter electrode (CHI127, CH Instruments). When using disk electrodes, the buffer was sparged with argon for 10 minutes immediately prior to scanning.

When using slide electrodes, no sparging was performed. CC was performed with the following parameters: a 3 s induction period at 0.2 V, a 0.25 s forward step period at -0.5 V, and a 1 s relaxation period at -0.5 V, with 125 sampling intervals. From 10 mM stock solution, Ruthenium (III) hexaammine ([Ru(NH_3_)_6_]^3+^, or RuHex) was added to the Tris-HCl until the buffer contained 50 μM RuHex. For disk electrodes, the solution was again sparged with argon for 10 minutes. For gold slide electrodes, the solution was pipetted up and down several times in the slide chamber and was allowed to sit between 30 seconds and 5 minutes. CC was again performed using the same parameters as before. All CC scans were performed three to four times in rapid sequence. Surface density was calculated using the assumption that one RuHex molecule binds to the anionic phosphodiester backbone of DNA for every three bases.^[19]^ Using the integrated Cottrell equation (**Equation 1**), it is possible to calculate the surface excess term.

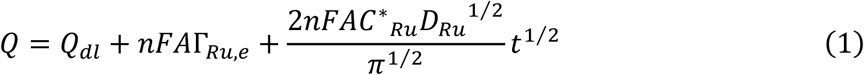

Where *n* is the number of electrons in the reaction (*n* = 1), *A* is the area of the working electrode (cm^2^), *F* is Faraday’s constant (C equiv^−1^), *D_Ru_* is the diffusion coefficient (cm^2^ s^−1^), *C\**is the bulk concentration (mol cm^−2^), *Q_dl_* is the double-layer, or capacitive charge (C), and *nFA*Γ*_Ru,e_* is the charge from the reduction of *Γ_Ru,e_* (mol cm^−2^) of adsorbed RuHex. By taking the *t =* 0 intercept of the Anson plots generated from the CC scans taken before and after the addition or RuHex, the values of *Q_dl_* and *Q* can be obtained, respectively. With *t =* 0, the diffusion component drops.

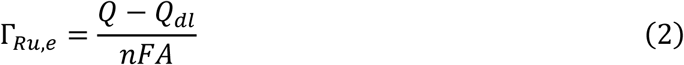

From here, the surface density of the dsDNA on the electrode (Γ*_dsDNA_*) can be calculated from the Γ*_Ru,e_* term using the relation

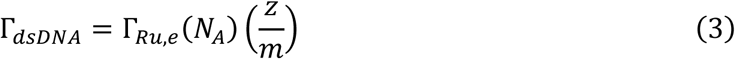

where *m* is the number of nucleotides in one DNA fragment (*m* = 1936), *z* is the charge of the redox molecule (*z* = 3), and *N_A_* is Avagadro’s number.

### Cell-Free Expression

PURExpress (E6800, New England Biolabs) and PUREfrex2.1 (PF213, GeneFrontier) cell-free protein synthesis kits were used for all TX-TL experiments. PURExpress is an all-in-one system and was used for many of the initial experiments. PUREfrex2.1 is a modular system and was used to express sfGFP in the absence of reducing agents. On-chip TX-TL reactions were carried out at room temperature directly on the surface of the DNA/MCH- modified electrodes for three hours.

For experiments where voltage is applied during CFE, an ITO-PET sheet is used as the counter electrode, separated by a thin silicone gasket (**Error! Reference source not found.**). When using this setup, only 7.5 μl of CFE reaction solution is used per 6 mm diameter monolayer area. For experiments where no voltage is applied, the ProPlate^®^ slide chambers remain on the slide, and either 20 μl or 10 μl CFE reaction solution onto each monolayer, depending on the surface area of the slide chambers to ensure total surface coverage. Slide chamber openings were covered with tape during the reaction to limit evaporation.

**Scheme 2.**
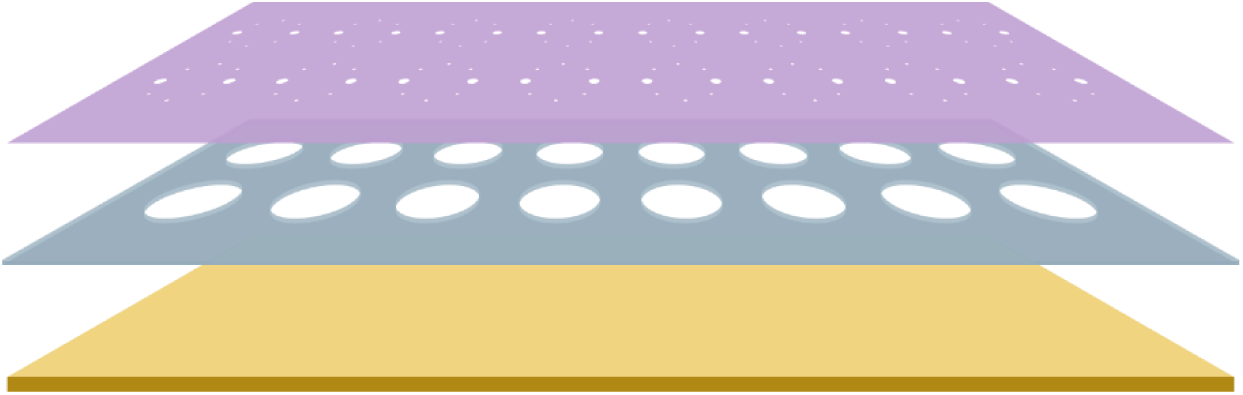
Reaction chamber setup for voltage application on planar gold slides. The DNA-functionalized gold surface (bottom) is connected to a potentiostat as the working electrode by adhesive copper tape. A 0.25 mm thick, clear silicone sheet with cut chamber holes aligned with the monolayer areas (center) is carefully placed onto the gold slide. Finally, an ITO-coated PET sheet with laser-cut portholes (top) is placed ITO-side down onto the silicone gasket. The ITO-PET sheet is connected to the potentiostat as the counter electrode by adhesive copper tape. CFE reaction solution is pipetted onto the surface through the portholes. These holes are then covered with tape or parafilm during the reaction to limit evaporation.

After on-chip incubation, a fraction of the cell-free reaction solution was aspirated and transferred to either the center of a Costar white transparent V-bottom 96-well storage plate (3363, Corning), or a Nunc transparent flat-bottom 384-well polystyrene plate (242757, Thermo Fisher). Fluorescence data for measuring sfGFP expression output or for fluorophore quantification was obtained using a BioTek Synergy HTX multi-mode plate reader with an excitation wavelength of 485±10 nm and an emission wavelength of 528±10 nm. Data from the plate reader was collected using Agilent Gen5 software.

### DNA Quantification

The DNA incubation concentrations (the concentration of DNA solutions immediately prior to immobilization) were measured using a NanoPhotometer (Implen). When attempting to measure DNA desorbed from the surface of an electrode, a Qubit 3.0 (Thermo Fisher) fluorometer running v1.02 of the instrument software was used with the Qubit dsDNA HS Assay Kit. Qubit readings falling below the assay’s limit of quantification were treated as a concentration of 0 ng/µL.

### Monolayer Decay Analysis

To evaluate the impacts both cell-free reagent exposure and storage under 1X PBS at 4 ℃ had on monolayer functionality, a linear mixed effect (LME) model was used. Coefficients were estimated via a random-intercept LME model fit using an expectation-maximization algorithm implemented in Python (v3.13.0; NumPy v2.4.2; SciPy v1.17.1).

Statistical significance of each fixed-effect coefficient was assessed using a two-sided t-test with conservative degrees of freedom (number of monolayers minus number of fixed effects); significance was defined a priori as α = 0.05, two-sided, for all statistical tests reported in this manuscript unless otherwise noted.

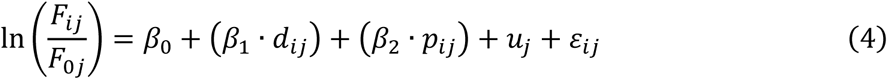

The response variable, ln(*F_ij_/F*_0*j*_), is the log-ratio of expression at measurement event *i* relative to the first measurement for monolayer *j*. *β*_0_ is the population-average intercept. The term (*β*_1_ · *d_ij_*) captures decay, with *d_ij_* representing the number of days between the previous TX-TL run and the current one, while (*β*_2_ · *p_ij_*) captures TX-TL exposure damage, with *p_ij_* representing the number of prior TX-TL runs already completed before measurement *i*. To account for inter-monolayer variability stemming from differences in surface quality, total packing densities, and heterogeneity of monolayer packing, each monolayer is assigned a random intercept of *u_j_*.

Finally, a residual capturing the intra-monolayer variation after the monolayer baseline *u_j_* has been accounted for is represented by *ε_ij_*. Log-ratio normalization was used prior to model fitting because dividing by F_0_ theoretically removes the monolayer-specific starting level, and converting to log scale addresses the assumption that decay is multiplicative rather than additive.

While this method was chosen for its simplicity, a notable caveat is that *F₀* is treated as a fixed reference rather than an estimated parameter, causing this ratio-based approximation to become less valid if initial measurement noise is large relative to the decay signal. To test key assumptions made by this normalization method, an alternative LME model with ln(*F_ij_*) as the response and ln(*F*_0*j*_) included as a covariate (not forced as a perfect offset with coefficient 1) was also tested (**Equation S1 and S2**, and **Table S2**).

### Fluorescence Intensity Measurements of DNA-Cy3 Layers on Gold

Once the 72-bp duplex was immobilized on 1000 Å gold slide electrodes, a transparent ITO top-plate was placed on top of the monolayer slide, separated by a 0.25 mm clear silicone gasket (identical to the setup shown in **Error! Reference source not found.**). 10 mM sodium chloride solution was added to the surface, and both the gold slide and ITO top-plate were wired up to a WaveNow XV potentiostat (Pine Research) running with AfterMath software using conductive copper tape. Fluorescence microscopy was performed on an OLYMPUS DP74 digital camera, mounted on an OLYMPUS BX53M TRF-S microscope with an X-Cite 120 LED Boost (Excelitas Technologies) light source set to 20% intensity. Videos were recorded using OLYMPUS Stream Essentials 2.2 software with a 4 s exposure time. A 5x objective lens (3.15x total magnification) was used. For processing, the mean grey value of each frame in the AVI files were extracted. Values from replicate monolayers where no potential was applied were used to subtract out the effects of photobleaching (**Figure S2**).

## 3 Results

### 3.1 Immobilization of gene-length dsDNA to a planar gold electrode surface

To begin developing our approach, we first immobilized double-stranded DNA fragments encoding the sfGFP gene with a thiol group attached to the promoter-proximal 5’ end. For initial experiments, standard, polished gold disk electrodes were used. We first investigated whether we could successfully attach a gene fragment of this size to the surface directly, and whether packing density can be controlled.

Packing density of DNA within the monolayer has considerable impact on the degree of conformational change we can induce through electrophoretic forces. If local crowding is high, the freedom of movement of any given strand within a cluster will be dependent on electrostatic and steric interactions with neighboring strands rather than on an applied surface charge alone. There are a variety of methods that can be used to control SAM density on an electrode surface, including co-adsorption with variable concentrations of alkanethiols such as MCH, adjusting salt concentrations during deposition,^[20]^ freezing the sample to increase density and uniformity,^[21]^ potential assisted deposition,^[22]^ and electrical desorption.^[15]^

To test whether dsDNA surface density could be controlled by varying the concentration of DNA in the bulk solution, we exposed six gold surfaces to either thiol-modified or unmodified 968 bp dsDNA at a range of concentrations. DNA bound to the surface was then quantified electrochemically, with technical triplicates averaged (**Figure 1**). We found that incubation concentration and final packing density were positively correlated, as expected. These numbers were further validated through alternative monolayer density quantification methods (**Methods S2**) where chemisorbed dsDNA was dehybridized and the resulting ssDNA was measured fluorometrically off-chip (**Figure S3**).

**Figure 1.**
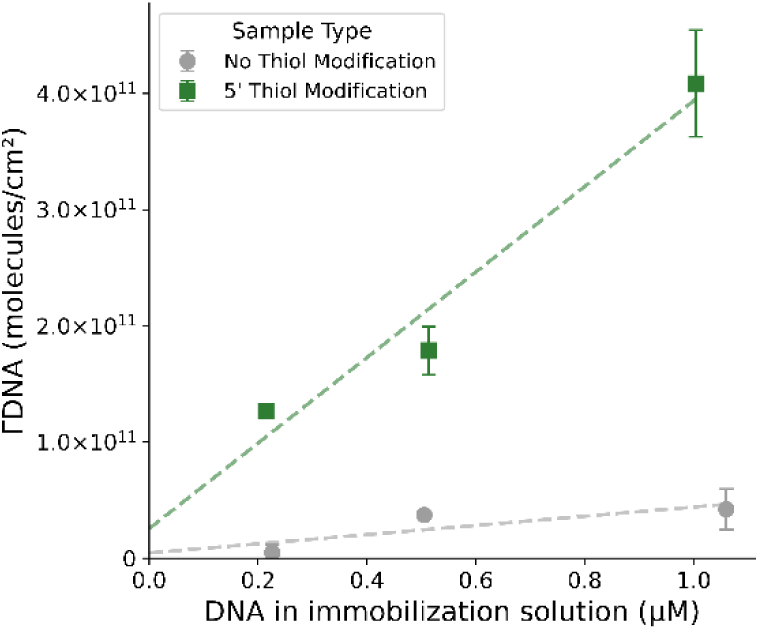
Density of dsDNA molecules immobilized on gold disk surface as a function of DNA concentration in the bulk solution. dsDNA SAMs are formed on polished gold disk electrodes using either thiolated DNA or unmodified DNA, and a range of starting incubation concentrations. Densities are measured using the chronocoulometric method.

Overall, our calculated packing densities closely agree with densities observed in other studies using similarly long (718 bp) thiol-modified dsDNA fragments (4.6 × 10^11^ molecules cm^−2^).^[9]^

### 3.2 Cell-Free TX-TL on Gold-Thiol-bound DNA SAMs

In order to conduct multiple attachment experiments in parallel, we constructed a multiplexable gasket array on patterned gold-coated microscope slides that allowed us to form and perform reactions on up to 16 spatially separate monolayer areas. CFE solution was applied to SAM areas. After incubation at room temperature, the CFE solution was extracted from the surface of the gold and sfGFP fluorescence was measured to evaluate expression efficiency.

#### 3.2.1 Re-use and storage conditions of monolayers

An important goal of our CFE platform is reusability. Therefore, we performed repeated CFE experiments with the same monolayers over the course of several days. All expression experiments were performed using commercially available cell-free reaction kits based on the PURE system. The first iterations of these re-use experiments were also performed under a constant applied voltage of either + 0.5 V or – 0.5 V to investigate whether a maintained, unidirectional electric field would impact transcription rates. Although initial exposure to cell-free TX-TL reagents often produced very clear expression signal, subsequent experiments using the same monolayers yielded significantly lower or fully deteriorated expression responses (**Figure 2**).

**Figure 2.**
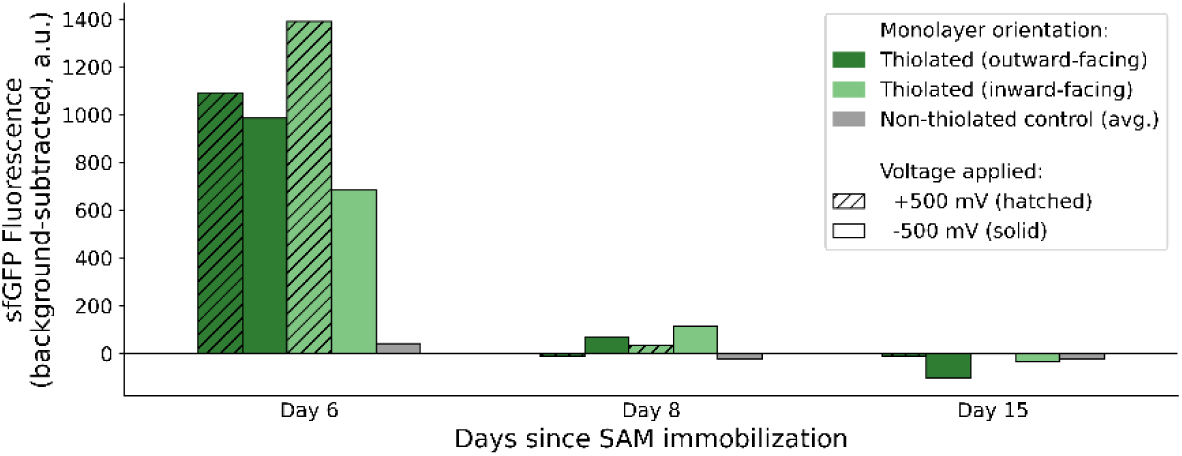
Cell-free sfGFP expression using DNA SAMs with applied charge over repeated uses. Six monolayers split evenly across two gold microscope slides were exposed to a constant applied potential of either + or – 500 mV (hatched and unhatched, respectively) throughout the duration of the 3 hour CFE reaction, with one gold microscope slide receiving the positive voltage, and the other receiving the negative voltage. Slides were randomly assigned to receive either the positive or negative applied potential prior to monolayer formation. Monolayers were formed using a 968 bp sfGFP gene, oriented so that the thiol group was attached to either the end closest to the promoter region (dark green), or furthest from the promoter region (light green). Mixed MCH-DNA monolayers were also formed using unmodified gene fragments (grey) to serve as wash controls.

Drops in expression efficiency also appeared to be independent of the time between the first and second experimental run. For many of our initial experiments, monolayers were stored dry at 4 ℃, under argon gas. This storage strategy was chosen given the known stability of dsDNA when stored dehydrated. While past investigations into storage outcomes of aptamer- based electrode sensors using similar gold-thiol binding chemistries have shown a roughly 75% aptamer signal loss when stored dry at ambient temperatures for more than a few days,^[23,24]^ storage at 4 ℃ should retard monolayer deterioration considerably.^[25,26]^ This assumption is supported by the observation of expression signal upon first exposure to cell-free reagents, even after the electrodes had been stored dry at 4 ℃ for multiple days (**Figure 2**). We also do not expect the act of drying the monolayers to lead to desorption or deterioration of DNA within the monolayers.^[24]^ The deterioration of monolayer functionality during or after first exposure to cell- free reagents, then, is most likely explainable by factors outside of storage conditions alone.

Previous studies have demonstrated how the rate of biofouling is partially dependent on the periodicity of electrochemical scanning, likely due to electric-field-induced desorption and rearrangement of the biofouled layer.^[26,27]^ The sustained voltage applied to the functionalized electrode surfaces during CFE, then, likely led to accelerated monolayer desorption. An additional factor worth considering was that any reducing agent in the cell-free TX-TL mix could cause the chemical decoupling of DNA from the gold. The majority of *E. coli*-based cell-free

TX-TL formulations include dithiothreitol (DTT) or other reducing agents such as 2- mercaptoethanol (2-ME) or tris(2-carboxyethyl)phosphine (TCEP) to replicate the native reducing environment within *E. coli*.^[28–30]^ Off-chip expression experiments using the same linear sfGFP gene to compare formulations with and without added DTT showed only minimal decrease in expression rate (**Figure S4**), however, lending viability to removal of DTT from our on-chip reactions.

#### 3.2.2 Expression signal decay is predictable after initial TX-TL exposure

To control any potential decoupling effects DTT might have on the monolayer, DTT was removed from the expression reaction system entirely. While average GFP fluorescence levels dropped upon the removal of DTT from the CFE reactions, signal remained well above background. Several DNA SAMs were fabricated using a range of starting DNA concentrations. Monolayers were stored in 1X PBS at 4 ℃ for less than 24 hours between immobilization and electrochemical interrogation, and for another roughly 72 hours before first exposure to TX-TL reaction mix. Fifteen thiolated DNA mixed monolayers, as well as one MCH monolayer and thirteen negative control mixed monolayers using unmodified DNA, were then exposed to DTT-free TX-TL reaction mix. End-point expression of sfGFP was plotted against measured monolayer density (**Figure 3A**). We observed high variability in endpoint CFE. The results fit weakly to a hill function (*R*^2^ = 0.39), with the majority of observed monolayer densities appearing to fall into an expression saturation regime where end-point expression is mostly independent of monolayer density.

**Figure 3.**
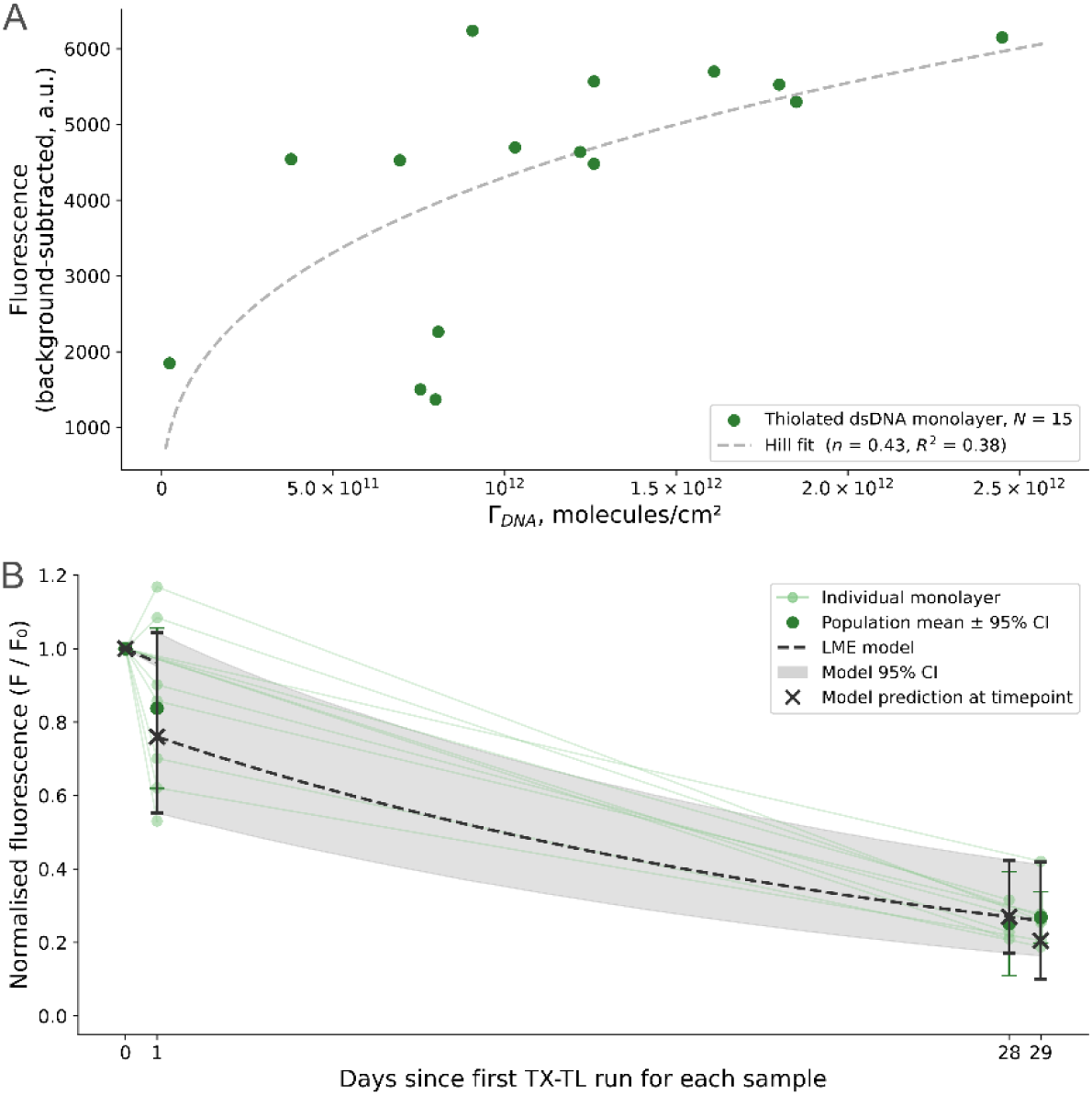
(**A**) Surface DNA density versus first exposure expression. Fifteen 968 bp dsDNA monolayers containing the sfGFP gene were fabricated on gold electrodes patterned on glass microscope slides using a range of incubation concentrations. The DNA was oriented with the promoter region situated proximal to the electrode surface. Densities were measured using the chronocoulometric method in open air. Cell-free TX-TL was carried out on the monolayers at room temperature and using a PURE-based system without any added DTT or other reducing agents for 3 hours. Some monolayers were exposed to a buffer solution representative of the PURE buffered environment (this buffer solution was not measured). Expression shows high variability and weakly fits a hill function, with the majority of observed monolayer densities appearing to fall into an expression saturation regime where end-point expression is mostly independent of monolayer density. (**B**) Signal retention (F/F0). After first exposure, SAMs were stored under PBS at 4 ℃. Cell-free TX-TL was carried out in the same manner as before the following day using a subset of the monolayers, including those that had only been exposed to buffered solution. A third and final TX-TL reaction was carried out 28 days later (while only three measurements were taken, four timepoints appear on Figure 3B because some monolayers were not exposed to TX-TL mix on the first day). An LME model was fit to the data to estimate the decay contributions attributable to storage and to TX-TL reagent exposure.

At the same time, three thiolated DNA mixed monolayers and two MCH monolayers were exposed to buffered solution formulated to imitate the buffer environment of PURE-based cell-free systems (50 mM pH 7.6 HEPES-KOH, 100 mM potassium glutamate, 13 mM magnesium acetate, 2 mM spermidine).^[31,32]^ After reaction solutions were collected and measured, all monolayers were rinsed at least twice in 10 mM Tris-HCl and then stored in 1X PBS at 4 ℃ overnight. The following day, on-chip TX-TL was repeated, this time with all monolayers including those that had been incubated in PURE analog buffer. Afterwards, monolayers were again stored in 1X PBS at 4 ℃, this time for 28 days before a third and final on-chip TX-TL run.

An LME model was designed to estimate the individual impacts of both cell-free reagent exposure and of storage duration on relative on-chip CFE levels (**Equation 4**). The response variable for the model was set as ln(*F_ij_*⁄*F*_0*j*_), the log-ratio of expressed sfGFP fluorescence measured after CFE event *i* for monolayer *j*, over the fluorescence measured after the very first CFE event for monolayer *j*. Model fitting on a sample set of 11 monolayers shows an estimated storage driven decay of 3.77% per day, and a TX-TL driven decay of 21.01% per exposure (**Figure 3B**). While it should be noted that only the storage attributable decay value is statistically significant (*p* = 0.0002 for storage decay versus *p* = 0.1817 for TX-TL decay), the approximately 21% estimated drop in expression after exposure to cell-free reagents aligns closely with an approximately 20% and 25% drop in nuclease-resistant aptamer signal observed by Leung et al. after exposure to DNAse I in PBS and to whole blood, respectively.^[33]^ Given that we expect insignificant DNAse activity when using a PURE-based cell-free system, our results taken in tandem with what has been observed by others suggests that the biofouling-induced loss in expression is caused predominately by the fouling layer obstructing access to the coding regions of the DNA rather than by desorption of DNA from the surface.

We adopted the ratio-normalized model, ln(*F_ij_*⁄*F*_0*j*_), as our primary model for estimating storage and TX-TL exposure decay rates, since it expresses decay as a single dimensionless fraction that is directly interpretable for experimental optimisation purposes. This model assumes that the proportional retention of signal is independent of the initial expression level *F*_0_. Put another way, it assumes that the coefficient relating ln(*F*_0_) to subsequent ln(*F*) is exactly 1. To test this assumption, we fit a secondary model in which this coefficient (γ) is estimated freely rather than constrained (**Equation S1 and S2**). The 95% confidence interval for γ included 1.0 (γ = 1.172, 95% CI [0.81, 1.53]), indicating that the independence assumption is not statistically rejected, and the decay rate estimates from both models agreed closely (storage: 3.77% vs. 3.72% per day; TXTL: 21.0% vs. 22.2% per exposure) (**Figure S5A, B**). We therefore report the log-ratio normalized model as our primary result, with the secondary relaxed model serving as a robustness check on its key normalization assumption. We note that the point estimate for γ was directionally above 1 (1.172), suggestive of higher-expressing monolayers retaining a slightly larger fraction of their signal over time; however, given that the 95% CI for this estimate spans 1.0, this directional trend should be treated as a hypothesis for future investigation with larger sample sizes rather than a confirmed effect.

#### 3.2.3 Biofouling-driven decay predominately stems from desorption of physisorbed molecules and steric hinderance of chemisorbed molecules

To interrogate decay driven by biofouling, sfGFP-encoding monolayers were formed with the promoter regions oriented at the proximal end relative to the gold electrode. Monolayers were also fabricated using unmodified DNA of the same sequence, and DNA-free (i.e. MCH only) monolayers were included to be used for background subtraction purposes. Fractions of these monolayers were exposed to one of two concentrations of bovine serum albumin (BSA), one of two concentrations of DTT, or a combination of BSA and DTT at concentrations believed to be approximately representative of the overall mass-per-volume protein concentrations and molar DTT concentrations commonly found in commercially available PURE expression systems. After approximately 2.5 hours of exposure, these solutions were collected from the monolayer surfaces and dsDNA was measured using fluorometric methods. Monolayers were washed twice with 1X PBS and then stored at 4 ℃ in fresh 1X PBS overnight before being removed and exposed to DTT-free TX-TL reaction mix for 3 hours.

The results show little to no detectible impact on either DNA desorption or on subsequent expression output after exposure solely to DTT. This was somewhat unexpected given our assumption that DTT, which is frequently used to destabilize disulfide bonds, would readily break the gold-thiol bonds tethering the chemisorbed DNA to the surface (**Figure S6A**). A Kruskal-Wallis test across all six exposure conditions indicated a significant overall difference in desorbed DNA for the thiol-modified monolayers (H = 18.536, p = 0.0023), and two-sided permutation tests (100,000 resamples) comparing each condition to buffer-only controls found no significant increase in desorption for either DTT concentration (p = 0.401 for 1 mM, p = 0.372 for 2 mM). However, exposure to BSA does appear to have a significant effect on monolayer desorption versus buffer-only exposure (**Figure S6B**) (permutation test yielded p = 0.028 or 0.029 across the three BSA-containing conditions). It should be noted however, that BSA on its own appears to be able to induce fluorescence of the dye used in the fluorometer assay. False positive results were observed when measuring blanks of BSA that had not been exposed to DNA monolayers (data not shown, but used for background subtraction). For samples containing both BSA and DNA, we proceeded with the expectation that the fluorescence attributable to BSA and to DNA individually to equal the total measured fluorescence, allowing us to get an accurate estimate of desorbed DNA after proper background subtraction.

Interestingly, desorption driven by BSA exposure appears to hold a strong positive correlation with subsequent end-point expression output only when the monolayer contains no thiolated DNA (**Figure 4A**). The amount of coding DNA remaining on the surface after the MCH incubation and washing steps included at the end of the SAM fabrication protocol is expected to be very small. It follows that at a low enough densities, the number of DNA molecules in the TX-TL reactions become rate limiting, and slight differences in the densities of immobilized strands across electrodes would be reflected in both the quantities of displaced DNA and in the end-point sfGFP fluorescence measurements. We do not see this same correlation between desorbed DNA quantities and end-point expression from the monolayers formed using thiolated DNA. With an expected density within the expression-saturation area seen in **Figure 3A**, we would not expect to see much, if any, statistically significant changes to expression levels if the remaining physisorbed fraction were displaced by the biofouling layer. This assumption aligns with what we see from the data (**Figure 4B**). This is consistent with a Kruskal-Wallis test showing no significant treatment effect on expression (H = 6.780, p = 0.2375). The conclusion that the biofouling layer primarily disrupts physisorbed molecules while having a comparatively minor effect on the desorption of chemisorbed molecules is further supported by the findings of Leung et al.^[33]^

**Figure 4.**
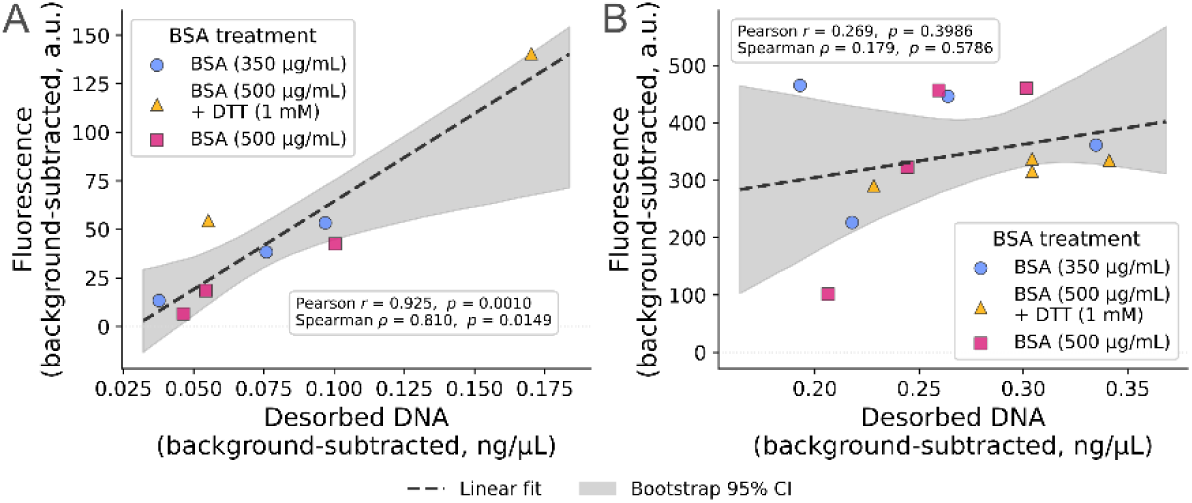
DNA desorbed from monolayers during 2.5 hour exposure to BSA or a combination of BSA and DTT, versus post-exposure TX-TL expression. Each datapoint represents an individual monolayer. All datapoints shown were from mixed monolayers composed of 968 bp sfGFP genes and an MCH backfilling layer. (**A**) Strong positive correlation in physisorbed monolayers, where all DNA strands were unmodified, and thus only a fraction of immobilized strands are expected remain on the surface after MCH backfilling (n = 8 monolayers, pooled across the three BSA-containing exposure conditions). (**B**) No correlation in chemisorbed monolayers, where all DNA strands were modified with a 5′ C6 thiol group at the proximal end, and thus were expected to be immobilized via gold-thiol bonds (n = 12 monolayers, pooled across the three BSA-containing exposure conditions).

A final observation worth noting is the small jump in the averaged desorbed DNA concentrations when SAMs are exposed to both BSA and DTT simultaneously. While the increase is likely not significant in a statistical sense (this specific comparison was not formally tested here), it may be suggestive of a combined effect where the DTT is able to decouple a small portion of the chemisorbed DNA, and then the BSA is able to physically displace the previously chemisorbed DNA before it has the chance to reform the gold-thiol bond.

### 3.3 Electrophoretic manipulation of relevant strand area

A unique benefit that comes with using a bio-functionalized electrode as a platform for enzymatic reactions is the ability to electrophoretically induce direction movement or conformational change of biological reaction components that are near or bound to the surface. This is especially true of a highly charged polyanion such as DNA. The ability to influence the height and orientation of both single- and double-stranded DNA monolayers by way of electric field has been understood for decades.^[34]^ It is also well understood that hybridization rates of aptameric sensors to target strands can be enhanced through the application of a positive surface potential, and even the reverse process of DNA duplex melting can be assisted electrochemically.^[11]^

We expect that for cell-free gene expression, the most critical portion of the gene in terms of regulatory control would be the promoter region. The T7 promoter region (TAA TAC GAC TCA CTA TAG G) in the sfGFP gene used in this study terminates 51 base pairs from the beginning of the fragment, and could be placed even further upstream in future studies. To demonstrate that dsDNA fragments can be electrophoretically manipulated up to or beyond a relevant height, potential-driven changes in strand conformation were identified by observing changes in the fluorescence of Cy3 fluorophores attached to the distal ends of gold-tethered, 72- bp duplex. The fluorescence strongly depends on the distance between the fluorophore and the gold due to non-radiative energy transfer to surface plasmons in the gold, which suppresses fluorescence the closer the dye is to the surface (**Figure 5A**).

**Figure 5.**
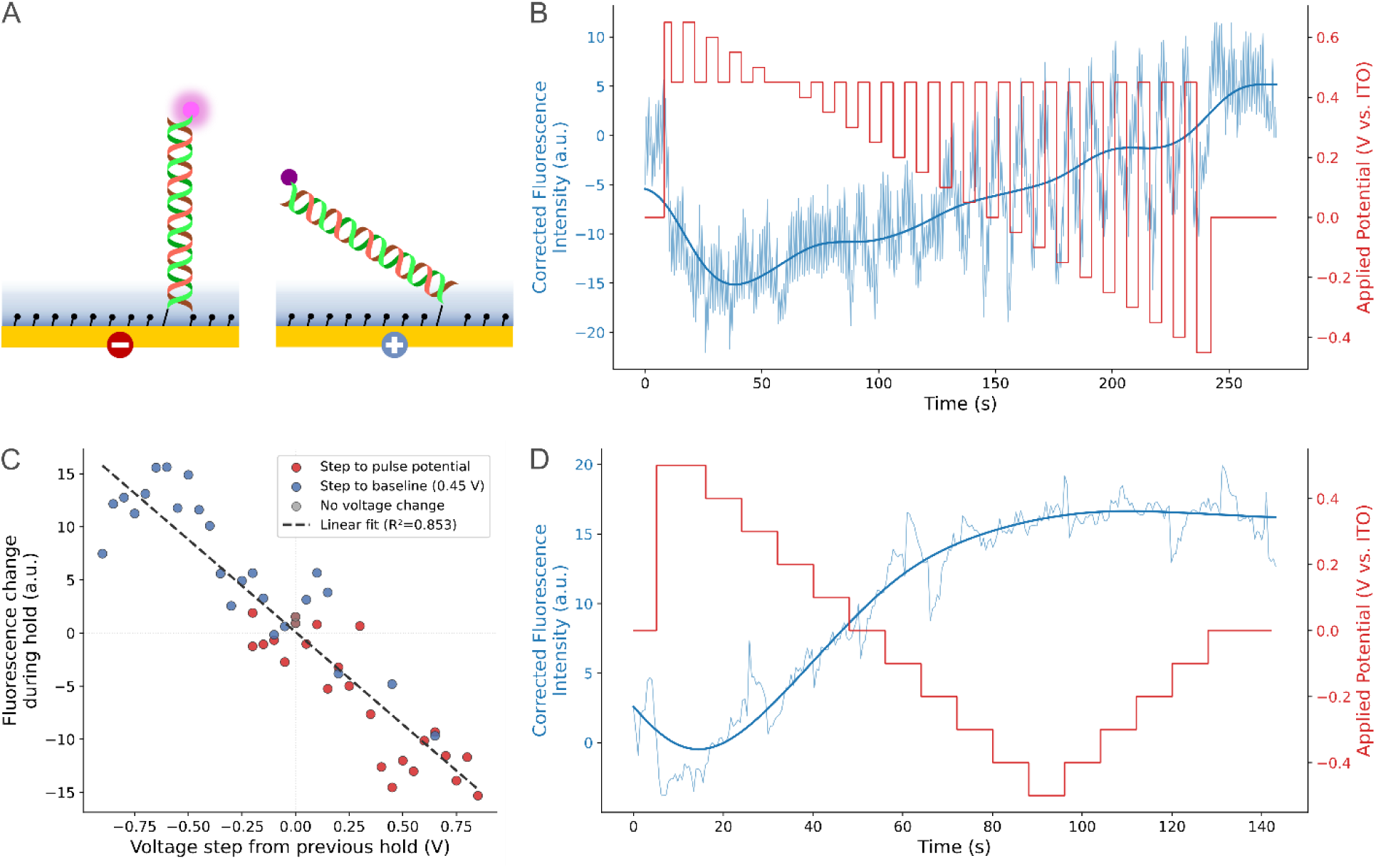
Tracking electrophoretically-driven conformational changes of dsDNA by measuring Cy3 fluorescence. (**A**) The applied electric field switches the negatively charged dsDNA fragment between an upright (left) and collapsed (right) conformation depending on the surface charge. A Cy3 fluorophore attached to the distal end of the DNA is quenched due to nonradiative energy transfer to surface plasmons in the gold, with the quenching effect becoming more dramatic the closer the Cy3 is from the electrode surface. (**B and D**) Applied potential (red solid line, right-hand y-axis) is shown mapped on top of Cy3 fluorescence (light-blue trace, left-hand y-axis) measured by taking the mean grey value of each frame of a fluorescence microscopy video. Cy3 fluorescence values have been corrected for photobleaching. The smoothed fluorescence (solid blue line) shows that as voltage decreases, the average distance of the DNA strands from the surface gets higher. (**B**) A switchable response is shown using a mixed monolayer of 72-mer fragments with a 5′ C6 thiol modification and a 3′ Cy3 modification, and the (**C**) fluorescence response versus voltage step magnitude. The fluorescence response is defined as the change in fluorescence from the start of a potential step to the end of the same step, and the resulting linear fit (dotted line) for the 72-mer monolayer has an R^2^ of 0.852. (**D**) A gradual stepping response is shown using a mixed monolayer of 968 bp dsDNA with a 5′ C6 thiol modification and a 5′ Cy3 modification. This monolayer was measured once (n = 1).

Voltage was switched within a range of 0.6 and -0.4 V, returning to 0.45 V for 5 seconds before stepping down another 0.05 V (**Figure 5B**). An initial large positive potential applied to the surface results in a noticeable drop in fluorescence as the strands go from a relaxed, disordered state to a uniformly collapsed state. As the voltage becomes more negative and the pulse heights become larger however, fluorescence levels begin to trend upwards and the fluorescence oscillations become more pronounced. Notably, after photobleaching effects are accounted for, we see signal is returned to original fluorescence levels at approximately 0 V, before continuing to increase as voltage sweeps into the negatives. To quantify this relationship, we regressed the change in fluorescence during each hold against the magnitude and direction of the preceding voltage step across all 44 voltage transitions recorded for this monolayer (**Figure 5C**), revealing a significant, strongly negative linear relationship (R² = 0.852, slope = -17.66 a.u. V^−1^, two-sided p = 5.30 × 10^-19^, N = 44), consistent with fluorescence increasing as voltage steps become more negative and decreasing as voltage steps become more positive. The monolayer used in Figure 5B and 5C was run twice on separate days under similar conditions, with the trace and fit shown reflecting one representative run (n = 1 monolayer). The trace and fit shown, as well as the fluorescence response versus voltage step, of the second run are shown in **Figure S7A** and **Figure S7B**, respectively.

We performed similar experiments using full 968 bp gene fragments with Cy3 tagging the distal ends, and a signal response was still observed during potential application (**Figure 5D**).

Interestingly, fluorescence still increases monotonically as applied potential decreases despite the contour length (*L*) being roughly 6.58 times the persistence length (*P*). For a polymer where *L* ≥ 3*P*, the approximate formula for the root-mean-square end-to-end distance of a simple worm-like chain is

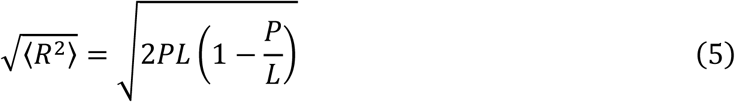

which gives us an estimated distal end height from the surface of ∼167 nm.^[35]^ However, given a buffer of 10 mM NaCl was used, and we expect the monolayers to be moderately to very densely packed (≥1.5·10^11^ molecules per cm^2^), the monolayers used in this experiment fall within the “salted brush” regime, as described by Bracha, et al.^[4]^ In this regime, the counterions that neutralize the negatively charged phosphate backbones are electrostatically confined within the brush volume, creating an ion concentration imbalance with the bulk solution that drives strand stretching. Electrostatic repulsion between the partially screened negatively charged backbones of densely packed neighboring strands also contributes to lateral pressure that favors a more upright orientation. In the salted brush

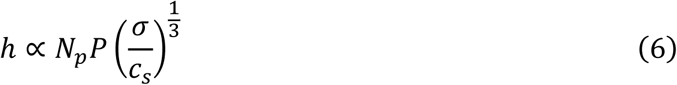

where *N_p_* is the number of persistence lengths, *σ* is the packing density, and *c_s_* is the salt concentration. Taking the power-law scaling of height with the ratio of density and ionic strength into account, it is reasonable to expect the average height of DNA strands within the monolayer at neutral potential to be slightly higher than the previously estimated ∼167 nm using the worm-like chain model, and for average height to increase with density. Due to this being beyond the region where energy transfer due to near-field coupling occurs and where damped interference oscillations begin to occur,^[15,36,37]^ we should be cautious to make relativistic comparisons between the fluorescence shifts observed with the 968 bp monolayers and the 72 bp monolayers. However, given the monotonic increase in the smoothed fluorescence as voltage becomes more negative, it is clear that the conformation of the bulk monolayer remains electrophoretically addressable even at full gene length. These findings suggest that we can reliably and reversibly manipulate the conformation of our genes up to and beyond the length that may be most relevant for on-chip CFE workflows.

Relevant to this study, the buffer components of a PURE-based protein expression system (50 mM HEPES-KOH, 100 mM potassium glutamate, 13 mM magnesium acetate, 2 mM spermidine) are estimated to have a total ionic strength around 170-190 mM, and therefore the Debye length of the solution on a gold electrode would only be ∼0.7 nm at ambient temperature.

Beyond this screening distance, the potential decays rapidly and Brownian motion begins to prevail over electric forces.^[15]^ This means the applied electric potential would effectively act only on the first two to three base pairs closest to the surface. It is not clear if this would be sufficient to induce conformational change up to the persistence length of ∼50 nm provided the monolayer is at a low enough density. The potential value at which the DNA conformation flips from an extended to a compact state (or vice versa) is referred to as the potential of conformation transition (*pct*) by Kaiser and Rant. They demonstrated that with increasing solution salinity, the attractive interaction between the negatively charged DNA polymer and its image charge within the metal surface is weakened. This means a more positive external potential is required to effectively pull the DNA towards the surface, meaning the *pct* has become more positive. In solutions containing very high concentrations of added salt, the abundant ions screen other charges with such efficiency that electrostatic potentials decay within sub-nanometer distances.^[15]^

## 4 Discussion

We have shown that room-temperature, cell-free gene expression using only the linearized genes immobilized to the surface of a gold electrode as the genetic template is possible and repeatable. We have also proposed a method for estimating the rate of loss of expression functionality due to reagent exposure and strand desorption during refrigerated storage under buffer. While the number of samples we used to derive estimated monolayer decay values was limited, thus effecting the statistical significance of the ∼21% per day decay value attributable to reagent exposure, this number aligns closely with an approximately 20 – 25% drop in nuclease-resistant aptamer signal observed in past studies investigating the impact of biofouling on similar DNA monolayers.^[33]^ The ∼3.8% per day decay value we attributed to monolayer decay during 4 ℃ storage in buffer is slightly higher than the < 3% aptamer loss over a 15 day period observed when functionalized gold nanoparticles are stored at 4 ℃ in buffer,^[25]^ but it is much closer to the ∼12.5% aptamer loss over a 7 day period observed when planar gold is stored at 4 ℃ in a more similar buffer.^[24]^ It should be noted, however, these past studies tracked strand desorption rather than expression signal loss, and while the two are likely linked, the relationship is not necessarily linear across all packing densities or strand lengths. Future work using gene monolayers on electrodes for repeated CFE may benefit from integrating a urea wash step after each expression experiment, since previous work has shown its effectiveness at recovering signal after biofouling.^[33,38]^ Adding a urea washing step is expected to significantly increase the lifetime of these chips.

Although we observed a sharp drop in on-chip CFE signal after applying a constant surface potential for the duration of the initial reaction, it is possible that an alternative, less intense voltage regime would have a much less significant impact on chip reusability. Applying surface potential in a pulsing pattern rather than as a constant potential could mitigate the effects of electric field-dependent monolayer deterioration and biofouling. In some instances, periodic negative excursions may help suppress thiol-to-disulfide oxidation by continually re-reducing any disulfide that forms.^[26]^ Further characterization of electrochemically driven monolayer desorption and biofouling should be performed as part of future biochip development to determine upper and lower voltage limits. These limits will likely be system and application specific, given SAM desorption may be dependent on thiol chain length,^[39,40]^ gold surface roughness,^[41]^ applied potential duration,^[33,42]^ and temperature,^[26]^ among other factors.

After intentionally fouling our monolayer chips with BSA and DTT, we observed a correlation between desorbed DNA quantities and post-exposure expression rates when the monolayer is sparse and entirely physisorbed, but not when it is dense and predominantly chemisorbed. These results are in line with the current understanding of how the biofouling layer forms within and on top of thiol-modified DNA mixed monolayers without substantially contributing to desorption of chemisorbed strands.^[33]^

Finally, we demonstrated that the conformation of dsDNA monolayers can be influenced by an applied electric field in a switchable and tunable manner. Our results using a 72mer duplex reflect those of past studies demonstrating rapid switching of sub-persistence length strands.^[15,34,43]^ This serves a baseline validation for electrophoretically mediated control of monolayer conformation. We were then able to demonstrate control over a 968 bp duplex using identical means. We originally did not expect to be able to exert any meaningful influence over conformation or monolayer height at that distance, given the rapid decay of potential just a few nanometers above the electrode surface and the worm-like chain behavior dsDNA or that length would normally exhibit. However, we noted a clear fluorescence signal responses attributable to the surface potential, which we posit is driven both by the applied potential experienced by the first few base pairs closest to the surface, as well as inter-strand repulsion along the rest of the gene fragment that helps the strands maintain some level of rigidity. These represent the initial steps taken towards the validation of DNA-functionalized planar electrodes as a modular platform for cell-free gene expression experiments.

One hurdle we encountered throughout on-chip TX-TL experimentation was a high degree of unpredictable variability in expression rates. While some of this variability is likely inherent to the cell-free reactions themselves,^[44]^ much of it likely stems from variability in electrode surface quality. This surface variability can be introduced during pre-immobilization cleaning and can lead to differences in surface area and in monolayer heterogeneity. This conclusion is based on our observations that there is typically more end-point variability between replicate monolayers when they are on different slides versus when they are on the same slide (data not shown). Future work may benefit from improved methods of both gold substrate preparation and surface quality control. Alternatively, using a methodology that allows for higher throughput at lower cost may help overcome the variability issue through the collection more datapoints. This may be explored by using a cheaper method of gold depositing, such as screen printing, or by reducing the size of the monolayer areas through microdroplet inkjet printing.

We envision DNA-functionalized electrodes as a new type of synthetic cell platform, and have therefore taken steps to characterize their robustness and electrochemical addressability. Tethering the genetic components of CFE reactions to an electronically addressable platform builds upon past work using silicon dioxide-bound DNA brushes^[3–6]^, which demonstrated novel passive expression control mechanisms that are dependent on brush height, density, composition, and orientation. Changing the binding chemistry from an electronically inert substrate to a planar electrode creates possibilities for novel functionalities and active control mechanisms including post-transcriptional aptameric RNA sensing,^[45,46]^ post-translational aptameric protein sensing,^[10,38,47,48]^ electrochemically driven hybridization,^[49]^ strand melting,^[11]^ strand desorption,^[50]^ and local pH control.^[12,13]^ If such a platform were to be designed as an array of electrodes rather than a single electrode, more functionalities could be imagined. For example, real-time spatial sensing could be used to track the speed of reaction cascades across the plane.

TX-TL of individual homogeneous gene patches could be turned “on” or “off” through electrochemical modulation of promoter-region hybridization/dehybridization or through electrophoretic modulation of steric hinderance. Finally, the gene-functionalized electrode could be integrated within a digital microfluidic device,^[51,52]^ allowing for active transport across the surface rather than relying on passive diffusion.

## Acknowledgements

Part of this work was conducted at the Washington Nanofabrication Facility as well as the Molecular Analysis Facility, which are National Nanotechnology Coordinated Infrastructure (NNCI) sites at the University of Washington supported in part by funds from the National Science Foundation (awards NNCI-1542101 and NNCI-2025489), the Molecular Engineering & Sciences Institute, and the Clean Energy Institute.

## Data Availability Statement

The data that support the findings of this study are available from the corresponding author upon reasonable request.

## References

1. A. C. Hunt, B. J. Rasor, K. Seki, et al., “Cell-Free Gene Expression: Methods and Applications,” Chemical Reviews 125, no. 1 (2025): 91–149. 10.1021/acs.chemrev.4c00116.

2. J. Cui, D. Wu, Q. Sun, et al., “A PEGDA/DNA Hybrid Hydrogel for Cell-Free Protein Synthesis,” Frontiers in Chemistry 8 (2020): 28. 10.3389/fchem.2020.00028.

3. A. Buxboim, M. Bar-Dagan, V. Frydman, D. Zbaida, M. Morpurgo, and R. Bar-Ziv, “A Single-Step Photolithographic Interface for Cell-Free Gene Expression and Active Biochips,” Small 3, no. 3 (2007): 500–510. 10.1002/smll.200600489.

4. D. Bracha, E. Karzbrun, S. S. Daube, and R. H. Bar-Ziv, “Emergent Properties of Dense DNA Phases toward Artificial Biosystems on a Surface,” Accounts of Chemical Research 47, no. 6 (2014): 1912–1921. 10.1021/ar5001428.

5. E. Karzbrun, A. M. Tayar, V. Noireaux, and R. H. Bar-Ziv, “Programmable On-Chip DNA Compartments as Artificial Cells,” Science 345, no. 6198 (2014): 829–832. 10.1126/science.1255550.

6. S. S. Daube, D. Bracha, A. Buxboim, and R. H. Bar-Ziv, “Compartmentalization by Directional Gene Expression,” Proceedings of the National Academy of Sciences 107, no. 7 (2010): 2836–2841. 10.1073/pnas.0908919107.

7. T. Thorsen, S. J. Maerkl, and S. R. Quake, “Microfluidic Large-Scale Integration,” Science 298, no. 5593 (2002): 580–584. 10.1126/science.1076996.

8. H. Niederholtmeyer, V. Stepanova, and S. J. Maerkl, “Implementation of Cell-Free Biological Networks at Steady State,” Proceedings of the National Academy of Sciences 110, no. 40 (2013): 15985–15990. 10.1073/pnas.1311166110.

9. M. Yang, H. C. M. Yau, and H. L. Chan, “Adsorption Kinetics and Ligand-Binding Properties of Thiol-Modified Double-Stranded DNA on a Gold Surface,” Langmuir 14, no. 21 (1998): 6121–6129. 10.1021/la980577i.

10. A. Langer, P. A. Hampel, W. Kaiser, et al., “Protein Analysis by Time-Resolved Measurements with an Electro-Switchable DNA Chip,” Nature Communications 4, no. 1 (2013): 2099. 10.1038/ncomms3099.

11. R. M. West, “Review—Electrical Manipulation of DNA Self-Assembled Monolayers: Electrochemical Melting of Surface-Bound DNA,” Journal of The Electrochemical Society 167, no. 3 (2020): 037544. 10.1149/1945-7111/ab67ad.

12. N. Fomina, C. A. Johnson, A. Maruniak, et al., “An Electrochemical Platform for Localized pH Control on Demand,” Lab on a Chip 16, no. 12 (2016): 2236–2244. 10.1039/C6LC00421K.

13. G. Pelossof, R. Tel-Vered, S. Shimron, and I. Willner, “Controlling Interfacial Electron Transfer and Electrocatalysis by pH- or Ion-Switchable DNA Monolayer-Modified Electrodes,” Chemical Science 4, no. 3 (2013): 1137. 10.1039/c2sc22193d.

14. J. H. Kim, J.-A. Hong, M. Yoon, M. Y. Yoon, H.-S. Jeong, and H. J. Hwang, “Solid-Phase Genetic Engineering with DNA Immobilized on a Gold Surface,” Journal of Biotechnology Sc, no. 3 (2002): 213–221. 10.1016/S0168-1656(02)00051-2.

15. W. Kaiser, and U. Rant, “Conformations of End-Tethered DNA Molecules on Gold Surfaces: Influences of Applied Electric Potential, Electrolyte Screening, and Temperature,” Journal of the American Chemical Society 132, no. 23 (2010): 7935–7945. 10.1021/ja908727d.

16. C.-Y. Lee, P. Gong, G. M. Harbers, D. W. Grainger, D. G. Castner, and L. J. Gamble, “Surface Coverage and Structure of Mixed DNA/Alkylthiol Monolayers on Gold: Characterization by XPS, NEXAFS, and Fluorescence Intensity Measurements,” Analytical chemistry 78, no. 10 (2006): 3326–3334. 10.1021/ac052137j.

17. J. Zhang, S. Song, L. Wang, D. Pan, and C. Fan, “A Gold Nanoparticle-Based Chronocoulometric DNA Sensor for Amplified Detection of DNA,” Nature Protocols 2, no. 11 (2007): 2888–2895. 10.1038/nprot.2007.419.

18. S. Moses, S. H. Brewer, L. B. Lowe, et al., “Characterization of Single- and Double- Stranded DNA on Gold Surfaces,” Langmuir 20, no. 25 (2004): 11134–11140. 10.1021/la0492815.

19. A. B. Steel, T. M. Herne, and M. J. Tarlov, “Electrochemical Quantitation of DNA Immobilized on Gold,” Analytical Chemistry 70, no. 22 (1998): 4670–4677. 10.1021/ac980037q.

20. K. Kobayashi, H. Tateishi-Karimata, K. Tsutsui, Y. Wada, and N. Sugimoto, “DNA Morphologic Changes Induced by Spermine on a Gold Surface under DNA Crowding Conditions,” Chemistry Letters 40, no. 8 (2011): 855–857. 10.1246/cl.2011.855.

21. Z. Li, Y. Lv, X. Duan, B. Liu, and Y. Zhao, “Highly Uniform DNA Monolayers Generated by Freezing-Directed Assembly on Gold Surfaces Enable Robust Electrochemical Sensing in Whole Blood,” Angewandte Chemie International Edition 62, no. 45 (2023). 10.1002/anie.202312975.

22. K. K. Leung, I. Martens, H.-Z. Yu, and D. Bizzotto, “Measuring and Controlling the Local Environment of Surface-Bound DNA in Self-Assembled Monolayers on Gold When Prepared Using Potential-Assisted Deposition,” Langmuir 36, no. 24 (2020): 6837– 6847. 10.1021/acs.langmuir.9b03970.

23. X. Zhou, J. Slaughter, S. Riki, C. C. Kuo, and A. Furst, “Polymer Coating for the Long-Term Storage of Immobilized DNA,” ACS Sensors 10, no. 7 (2025): 5019–5026. 10.1021/acssensors.5c00937.

24. J. Chung, A. Billante, C. Flatebo, et al., “Effects of Storage Conditions on the Performance of an Electrochemical Aptamer-Based Sensor,” Sensors & Diagnostics 3, no. 6 (2024): 1044–1050. 10.1039/D4SD00066H.

25. N. Bhatt, P.-J. J. Huang, N. Dave, and J. Liu, “Dissociation and Degradation of Thiol-Modified DNA on Gold Nanoparticles in Aqueous and Organic Solvents,” Langmuir 27, no. 10 (2011): 6132–6137. 10.1021/la200241d.

26. Z. Watkins, A. Karajic, T. Young, R. White, and J. Heikenfeld, “Week-Long Operation of Electrochemical Aptamer Sensors: New Insights into Self-Assembled Monolayer Degradation Mechanisms and Solutions for Stability in Serum at Body Temperature,” ACS Sensors 8, no. 3 (2023): 1119–1131. 10.1021/acssensors.2c02403.

27. A. Shaver, and N. Arroyo-Currás, “The Challenge of Long-Term Stability for Nucleic Acid-Based Electrochemical Sensors,” Current Opinion in Electrochemistry 32 (2022): 100902. 10.1016/j.coelec.2021.100902.

28. A. Zemella, L. Thoring, C. Hoffmeister, and S. Kubick, “Cell-Free Protein Synthesis: Pros and Cons of Prokaryotic and Eukaryotic Systems,” ChemBioChem 16, no. 17 (2015): 2420–2431. 10.1002/cbic.201500340.

29. D. Garenne, M. C. Haines, E. F. Romantseva, P. Freemont, E. A. Strychalski, and V. Noireaux, “Cell-Free Gene Expression,” Nature Reviews Methods Primers 1, no. 1 (2021): 49. 10.1038/s43586-021-00046-x.

30. D. Kim, and J. R. Swartz, “Efficient Production of a Bioactive, Multiple Disulfide-bonded Protein Using Modified Extracts of *Escherichia Coli*,” Biotechnology and Bioengineering 85, no. 2 (2004): 122–129. 10.1002/bit.10865.

31. Y. Shimizu, T. Kanamori, and T. Ueda, “Protein Synthesis by Pure Translation Systems,” Methods 36, no. 3 (2005): 299–304. 10.1016/j.ymeth.2005.04.006.

32. Y. Kuruma, and T. Ueda, “The PURE System for the Cell-Free Synthesis of Membrane Proteins,” Nature Protocols 10, no. 9 (2015): 1328–1344. 10.1038/nprot.2015.082.

33. K. K. Leung, A. M. Downs, G. Ortega, M. Kurnik, and K. W. Plaxco, “Elucidating the Mechanisms Underlying the Signal Drift of Electrochemical Aptamer-Based Sensors in Whole Blood,” ACS Sensors 6, no. 9 (2021): 3340–3347. 10.1021/acssensors.1c01183.

34. S. O. Kelley, J. K. Barton, N. M. Jackson, et al., “Orienting DNA Helices on Gold Using Applied Electric Fields,” Langmuir 14, no. 24 (1998): 6781–6784. 10.1021/la980874n.

35. C. Rivetti, C. Walker, and C. Bustamante, “Polymer Chain Statistics and Conformational Analysis of DNA Molecules with Bends or Sections of Different Flexibility,” Journal of Molecular Biology 280, no. 1 (1998): 41–59. 10.1006/jmbi.1998.1830.

36. J. R. Lakowicz, “Radiative Decay Engineering 5: Metal-Enhanced Fluorescence and Plasmon Emission,” Analytical Biochemistry 337, no. 2 (2005): 171–194. 10.1016/j.ab.2004.11.026.

37. M. Gómez-Castaño, A. Redondo-Cubero, L. Buisson, et al., “Energy Transfer and Interference by Collective Electromagnetic Coupling,” Nano Letters 19, no. 8 (2019): 5790–5795. 10.1021/acs.nanolett.9b02521.

38. A. Langer, M. Schräml, R. Strasser, et al., “Polymerase/DNA Interactions and Enzymatic Activity: Multi-Parameter Analysis with Electro-Switchable Biosurfaces,” Scientific Reports 5, no. 1 (2015): 12066. 10.1038/srep12066.

39. D. Oyamatsu, T. Fujita, S. Arimoto, H. Munakata, H. Matsumoto, and S. Kuwabata, “Electrochemical Desorption of a Self-Assembled Monolayer of Alkanethiol in Ionic Liquids,” Journal of Electroanalytical Chemistry 615, no. 2 (2008): 110–116. 10.1016/j.jelechem.2007.12.003.

40. T. Ghaly, B. E. Wildt, and P. C. Searson, “Electrochemical Release of Fluorescently Labeled Thiols from Patterned Gold Surfaces,” Langmuir 26, no. 3 (2010): 1420–1423. 10.1021/la9032282.

41. E. Pensa, C. Vericat, D. Grumelli, et al., “New Insight into the Electrochemical Desorption of Alkanethiol SAMs on Gold,” Physical Chemistry Chemical Physics 14, no. 35 (2012): 12355. 10.1039/c2cp41291h.

42. G. Sánchez-Pomales, L. Santiago-Rodríguez, N. E. Rivera-Vélez, and C. R. Cabrera, “Control of DNA Self-Assembled Monolayers Surface Coverage by Electrochemical Desorption,” Journal of Electroanalytical Chemistry 611, no. 1-2 (2007): 80–86. 10.1016/j.jelechem.2007.08.003.

43. U. Rant, K. Arinaga, S. Scherer, et al., “Switchable DNA Interfaces for the Highly Sensitive Detection of Label-Free DNA Targets,” Proceedings of the National Academy of Sciences 104, no. 44 (2007): 17364–17369. 10.1073/pnas.0703974104.

44. K. A. Rhea, N. D. McDonald, S. D. Cole, V. Noireaux, M. W. Lux, and P. E. Buckley, “Variability in Cell-Free Expression Reactions Can Impact Qualitative Genetic Circuit Characterization,” Synthetic Biology 7, no. 1 (2022): ysac011. 10.1093/synbio/ysac011.

45. Y. Yan, S. Ding, D. Zhao, R. Yuan, Y. Zhang, and W. Cheng, “Direct Ultrasensitive Electrochemical Biosensing of Pathogenic DNA Using Homogeneous Target-Initiated Transcription Amplification,” Scientific Reports 6, no. 1 (2016): 18810. 10.1038/srep18810.

46. Y.-H. Cheng, S.-J. Liu, and J.-H. Jiang, “Enzyme-Free Electrochemical Biosensor Based on Amplification of Proximity-Dependent Surface Hybridization Chain Reaction for Ultrasensitive mRNA Detection,” Talanta 222 (2021): 121536. 10.1016/j.talanta.2020.121536.

47. A. A. Rowe, R. J. White, A. J. Bonham, and K. W. Plaxco, “Fabrication of Electrochemical-DNA Biosensors for the Reagentless Detection of Nucleic Acids, Proteins and Small Molecules,” Journal of Visualized Experiments no. 52 (2011): 2922. 10.3791/2922-v.

48. F. V. Oberhaus, D. Frense, and D. Beckmann, “Immobilization Techniques for Aptamers on Gold Electrodes for the Electrochemical Detection of Proteins: A Review,” Biosensors 10, no. 5 (2020): 45. 10.3390/bios10050045.

49. T. Zhai, C. Sun, D. Ye, et al., “Electrochemically Driven Assembly of Framework Nucleic Acids,” Journal of Electroanalytical Chemistry 905 (2022): 115901. 10.1016/j.jelechem.2021.115901.

50. K. Sun, B. Jiang, and X. Jiang, “Electrochemical Desorption of Self-Assembled Monolayers and Its Applications in Surface Chemistry and Cell Biology,” Journal of Electroanalytical Chemistry 656, no. 1–2 (2011): 223–230. 10.1016/j.jelechem.2010.11.008.

51. L. Malic, T. Veres, and M. Tabrizian, “Biochip Functionalization Using Electrowetting- on-Dielectric Digital Microfluidics for Surface Plasmon Resonance Imaging Detection of DNA Hybridization,” Biosensors and Bioelectronics 24, no. 7 (2009): 2218–2224. 10.1016/j.bios.2008.11.031.

52. L. Malic, T. Veres, and M. Tabrizian, “Nanostructured Digital Microfluidics for Enhanced Surface Plasmon Resonance Imaging,” Biosensors and Bioelectronics 26, no. 5 (2011): 2053–2059. 10.1016/j.bios.2010.09.001.

