## Supplement for "Characterizing the Assembly and Functional Properties of Gene-Length Mixed DNA Monolayers on Electrodes for Cell-Free Expression"

**Supplementary Figures**

Figure S1: XRD spectrum of electron beam evaporated gold surface

Figure S2: Fluorescence trace of neutral voltage control monolayer used for photobleaching correction

Figure S3: Monolayer packing density measured by way of dehybridization

Figure S4: Fluorescence measurements of off-chip cell-free expression reactions using various formulations

Figure S5: Comparison of estimated expression decay values across two LME model variants

Figure S6: Impacts of BSA and/or DTT exposure on monolayer desorption and expression efficiency

Figure S7: Tracking electrophoretically-driven conformational changes of dsDNA by measuring Cy3 fluorescence

**Supplementary Tables**

Table S1: sfGFP gene sequence

Table S2: Comparison of model statistics

**Supplementary Methods**

Methods S1: Alternative LME model with relaxed day-one signal coefficient

Methods S2: Monolayer packing density quantification of 968 bp dsDNA strands via chemical stand melting

**Supplementary References**

### Supplementary Figures

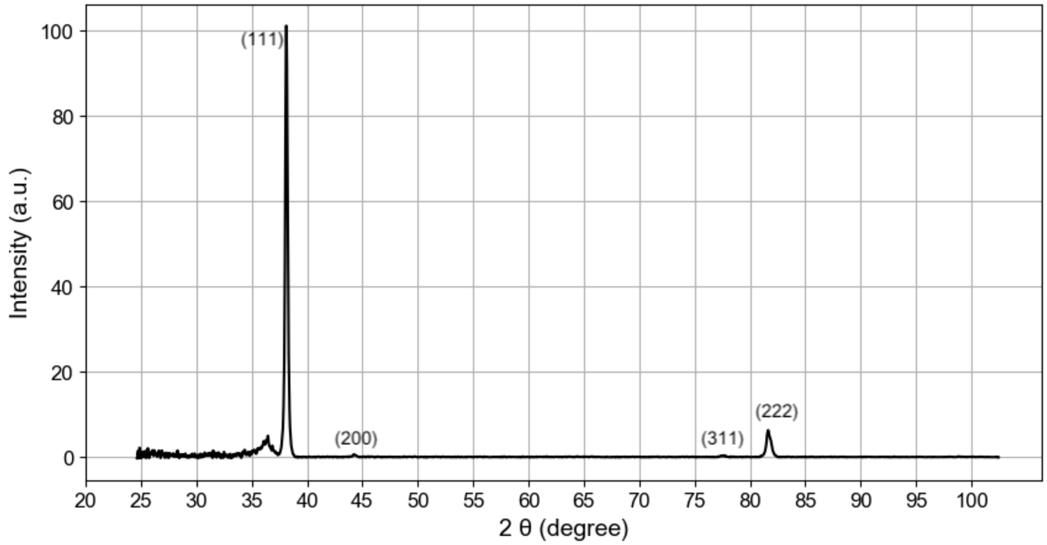


Figure S1: XRD spectrum of electron beam evaporated gold surface

Background subtracted XRD spectrum of an e-beam evaporated gold (1000 Å gold layer, with 50 Å chromium adhesion layer) microscope slide that had been used in previous SAM experiments indicating a predominantly Au<111> crystal structure. Presence of additional crystal structure peaks are labeled. The day prior to XRD analysis, the slide was cleaned in 60˚C Nano-Strip for 10 minutes and then O2 plasma cleaned for 5 minutes before being stored dry at room temperature.

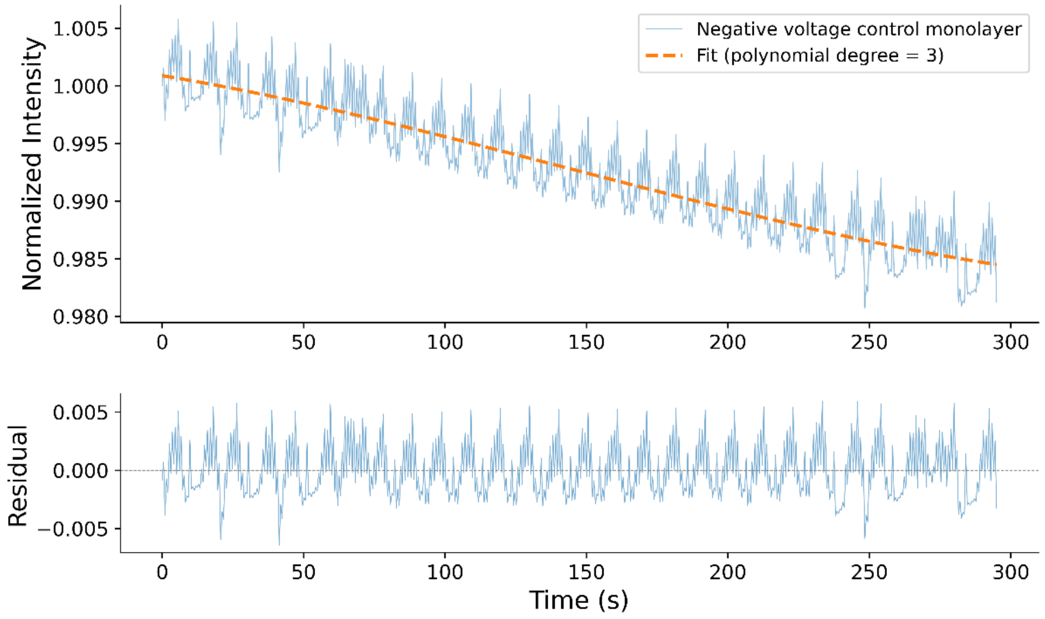


<u>Figure S2: Fluorescence trace of neutral voltage control monolayer used for photobleaching correction</u>

A 72 bp dsDNA fragment modified with a 5’ thiol group at one end and a 3’ Cy3 fluorophore at the other was used to form a monolayer on planar gold. A video was taken using a fluorescence microscope while the monolayer was under neutral voltage (i.e. the gold electrode was not connected to the potentiostat). Raw mean grey value data extracted from each frame of the AVI file were plotted (top, blue solid line), and a cubic trendline was generated to map the decrease in fluorescence attributable to photobleaching (top, orange dashed line). While a linear regression line would have also been sufficient for the data shown, a third-degree polynomial fit was selected because it was observed to fit the photobleaching effect seen in the 968 bp monolayers (data not shown) more cleanly. The photobleaching trend was subtracted from the original no-voltage control (bottom) as well as the voltage-exposed sample (**Figure 5B**).

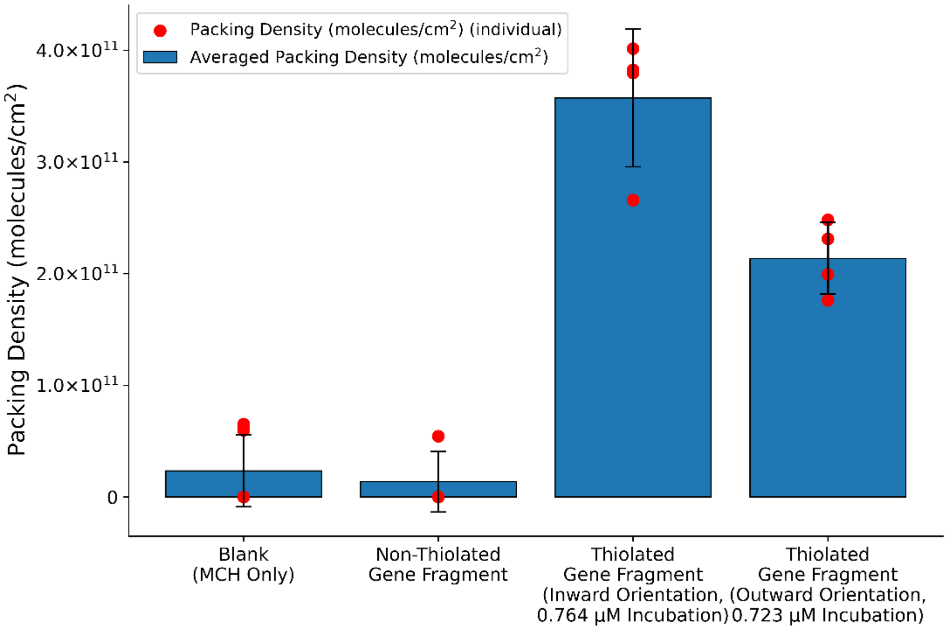


Figure S3: Monolayer packing density measured by way of dehybridization

Monolayers were formed and data was collected as described in Methods S2. Incubation concentrations of the non-thiolated gene variant, the inwardly oriented thiolated gene variant, and the outwardly facing thiolated gene variant were measured as 0.796 µM, 0.723 µM, and 0.764 µM, respectively. The dsDNA within the monolayers was dehybridized via exposure to 1 M NaOH. The DNA content of the NaOH post-exposure was measured fluorometrically. The measured packing densities all fall within the expected values given the measured incubation concentrations (**Figure 1**). Interestingly, there was an observable difference between the densities of the inward facing and outward facing strand orientations. This difference can at least in part be attributed to slight differences in initial incubation concentrations, however the effects of sequence-dependent conformational differences cannot be entirely ruled out

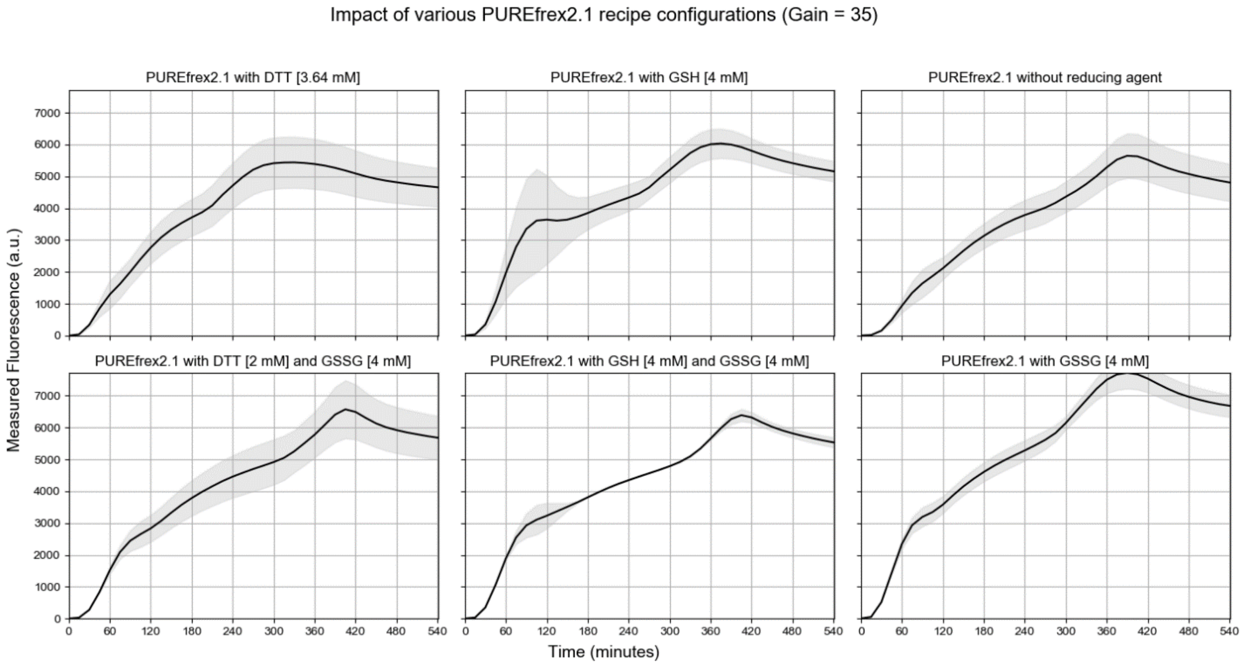


<u>Figure S4: Fluorescence measurements of off-chip cell-free expression reactions using various formulations</u>

PUREfrex2.1 cell-free expression mix with no added DNA was used for background subtraction. 4.5 µL duplicates of the six different formulations were prepared and aliquoted into v-bottom wells of a 96-well plate. Each formulation contained 25% v/v Solution I, 5% v/v Solution II, 10% v/v Solution III, 0.3 mM cysteine, and 2.5 ng/µL unmodified sfGFP gene fragment. An exception is the formulation with DTT as the sole additive (top-left), which, due to a dilution error, contained 22.73% v/v Solution I, 4.545% v/v Solution II, 9.09% v/v Solution III, 0.545 mM cysteine, and 2.27 ng/µL unmodified sfGFP gene fragment. Fluorescence measurements were taken every 15 minutes over the course of 9 hours at a constant setpoint of 29˚ C.

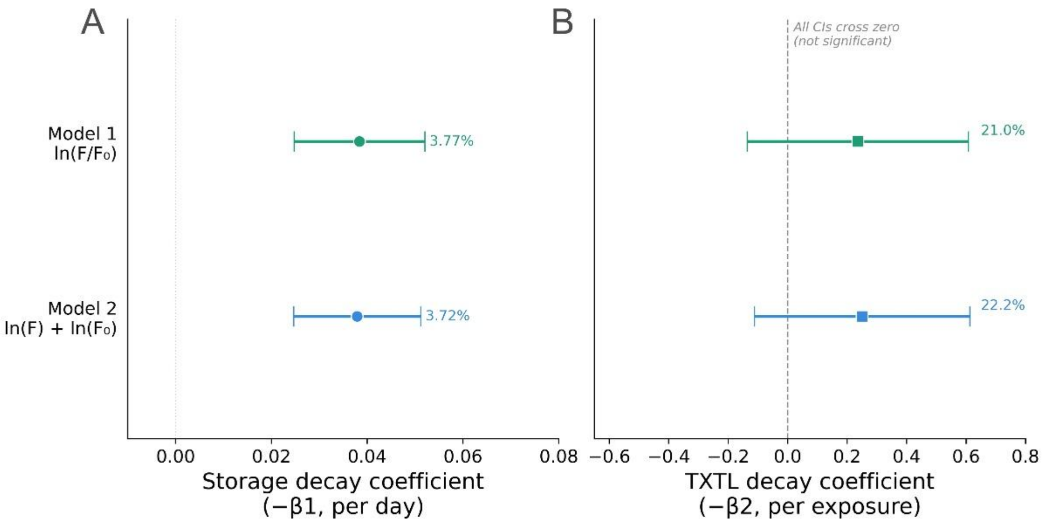


Figure S5: Comparison of estimated expression decay values across two LME model variants

Comparison of estimated percentage decay in end-point fluorescence of expressed sfGFP after 3 hours of cell-free TX-TL across two LME models. Model 1 is defined by Equation 4 in the main text, and Model 2 is defined by Equation S2 (**Methods S1**). PURE-based cell-free system was used with all DTT removed. Both (**A**) estimated per-day decay experienced by the SAM when stored in 1X PBS at 4 ℃, and (**B**) estimated per exposure decay experienced by the SAM after a 3 hour on-chip TX-TL run are comparable.

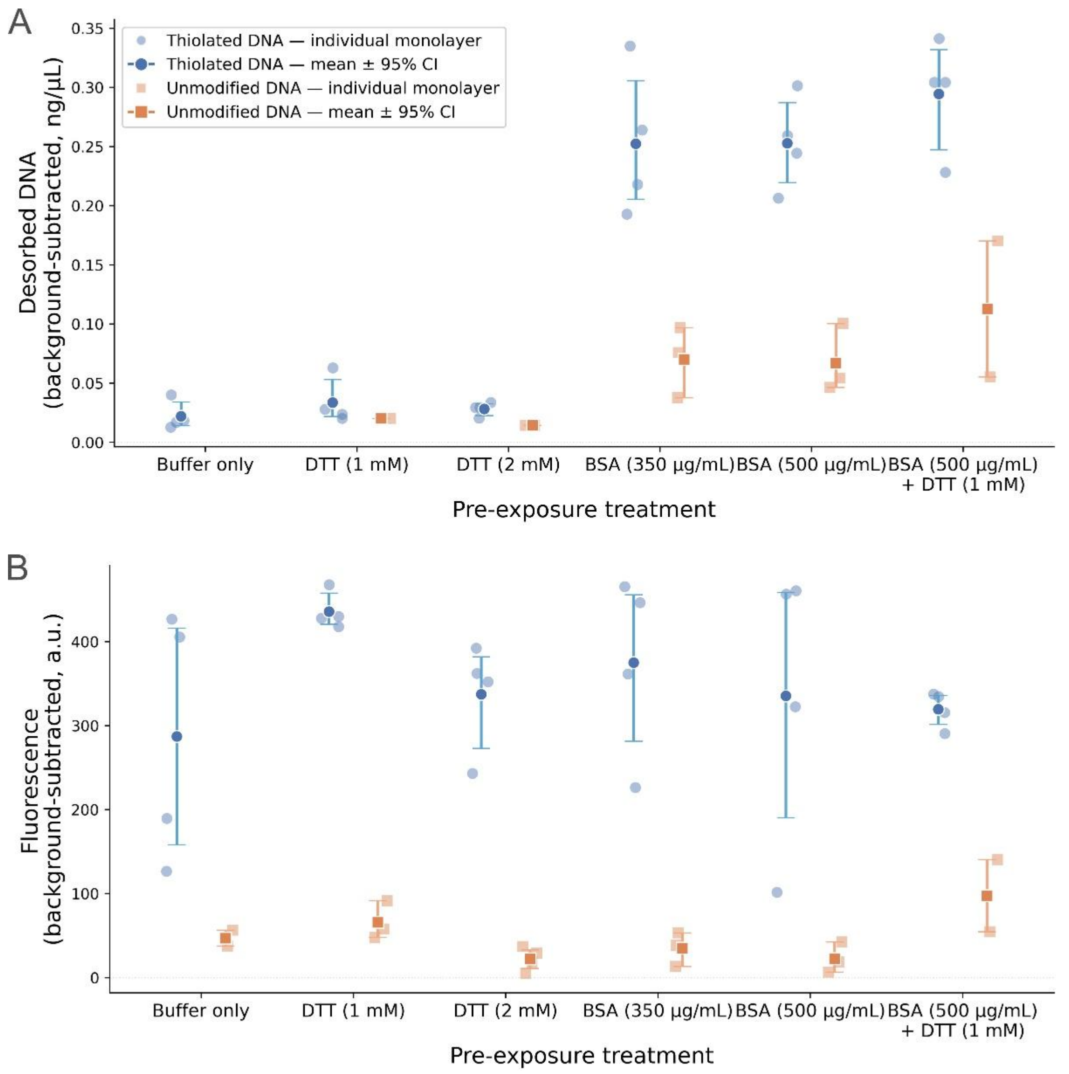


Figure S6: Impacts of BSA and/or DTT exposure on monolayer desorption and expression efficiency

Monolayers were exposed to buffered solution, DTT, BSA, or a combination of DTT and BSA. (**A**) After 2.5 hours of exposure at room temperature, the concentration of dsDNA in each of the exposure solutions was measured using a Qubit fluorometer. Solutions that were exposed to MCH-only monolayers were used for background subtraction. (**B**) Monolayers were rinsed with buffer, then exposed to DTT-free CFE solution for 3 hours at room temperature. After incubation, aliquots were taken and fluorescence was measured. Monolayers were formed in the following numbers per exposure condition: 3 blank (no DNA), 3 unmodified-gene, and 4 thiol-modified monolayers for each of 1 mM DTT and 350 µg/mL BSA; 4 blank, 4 unmodified-gene, and 4 thiol-modified monolayers for each of 2 mM DTT and 500 µg/mL BSA; and 2 blank, 2 unmodified-gene, and 4 thiol-modified monolayers for each of 500 µg/mL BSA + 1 mM DTT and buffer-only (60 monolayers total). One monolayer (unmodified gene, 500 µg/mL BSA exposure, no DTT) was excluded from the dataset (**Figure 4, Figure S6**) as an extreme outlier in desorbed DNA relative to the other replicate monolayers of the same type and exposure condition (background-subtracted desorbed DNA of 0.390 ng/µL, versus 0.046–0.100 ng/µL for the remaining replicates); its background-subtracted expression signal (5.5 a.u.) fell within the range observed for that treatment group (5.5–42.5 a.u.), indicating the sample was not anomalous with respect to expression.

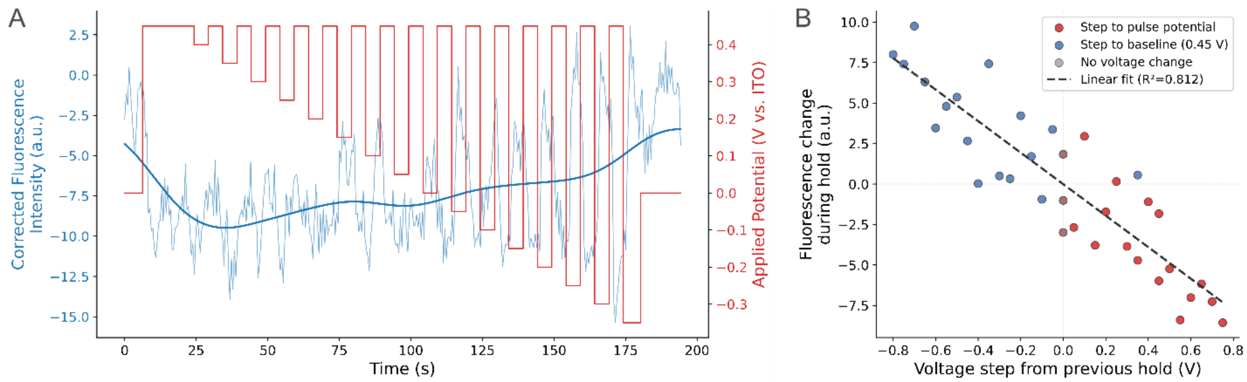


<u>Figure S7: Tracking electrophoretically-driven conformational changes of dsDNA by measuring Cy3 fluorescence</u>

(A) Applied potential (red solid line, right-hand y-axis) is shown mapped on top of Cy3 fluorescence (light-blue trace, left-hand y-axis) measured by taking the mean grey value of each frame of a fluorescence microscopy video. Cy3 fluorescence values have been corrected for photobleaching. The smoothed fluorescence (solid blue line) shows that as voltage decreases, the average distance of the DNA strands from the surface gets higher. A switchable response is shown using a mixed monolayer of 72-mer fragments with a 5` C6 thiol modification and a 3` Cy3 modification, and the (B) fluorescence response versus voltage step magnitude. The fluorescence response is defined as the change in fluorescence from the start of a potential step to the end of the same step. The monolayer used is the same as the one shown in Figure 5B and 5C, but the data shown here was generated on a separate day.

**Supplementary Tables**

Table S1: sfGFP gene sequence

The below sequence shows the sense strand, read from 5` to 3`, of the 968 bp sfGFP gene duplex used throughout this text.

T7 Promoter

sfGFP coding sequence

T7 Terminator

ATCTCCAGTGGTTCAGGTACTCCCGCGAAATTAATACGACTCACTATAGGGAGACCA
CAACGGCCCCTGGGTACAGGGGTAAATAAAGTTATTGAGATAAGGAGGTATTTTATG
AGCAAAGGAGAAGAACTTTTCACTGGAGTTGTCCCAATTCTTGTTGAATTAGATGGT
GATGTTAATGGGCACAAATTTTCTGTCCGTGGAGAGGGTGAAGGTGATGCTACAAA
CGGAAAACTCACCCTTAAATTTATTTGCACTACTGGAAAACTACCTGTTCCATGGCC
AACACTTGTCACTACTCTGACCTATGGTGTTCAATGCTTTTCCCGTTATCCGGATCAC
ATGAAACGGCATGACTTTTTCAAGAGTGCCATGCCCGAAGGTTATGTACAGGAACG
CACTATATCTTTCAAAGATGACGGGACCTACAAGACGCGTGCTGAAGTCAAGTTTGA
AGGTGATACCCTTGTTAATCGTATCGAGTTAAAAGGTATTGATTTTAAAGAAGATGGA
AACATTCTCGGACACAAACTCGAGTACAACTTTAACTCACACAATGTATACATCACG
GCAGACAAACAAAAGAATGGAATCAAAGCTAACTTCAAAATTCGCCACAACGTTGAA
GATGGTTCCGTTCAACTAGCAGACCATTATCAACAAAATACTCCAATTGGCGATGGC
CCTGTCCTTTTACCAGACAACCATTACCTGTCGACACAATCTGTCCTTTCGAAAGAT
CCCAACGAAAAGCGTGACCACATGGTCCTTCTTGAGTTTGTAACTGCTGCTGGGAT
TACACATGGCATGGATGAGCTCTACAAATGATGAAAGGGTGATCCGGCTGCTAACAA
AGCCCGAAAGGAAGCTGAGTTGGCTGCTGCCACCGCTGAGCAATAACTAGCATAAC
CCCTTGGGGCCTCTAAACGGGTCTTGAGGGGTTTTTTGGCCGTTGTTTGTACTGTG
AT

Table S2: Comparison of model statistics

| | Model 1 | Model 2 |
|---|---|---|
| Storage-linked decay (% / day) * | 3.77 | 3.72 |
| TX-TL-linked decay (% / day) * | 21.0 | 22.2 |
| ln(F0) coefficient | 1.0 (fixed) | 1.172 |
| ICC | 0.848 | 0.825 |
| $\sigma^2_u$ | 0.05619 | 0.04870 |
| $\sigma^2_\varepsilon$ | 0.01005 | 0.01032 |
| $R^2$ marginal | 0.845 | 0.927 |
| $R^2$ conditional | 0.989 | 0.993 |
| Log-likelihood | 2.765 | 3.357 |
| AIC | 4.470 | 5.286 |
| BIC | 8.636 | 10.285 |

* Calculated using transformation for exponential decay, $(1-\exp(|\beta|))\times100$

**Supplementary Methods**

Methods S1: Alternative LME model with relaxed day-one signal coefficient

$$\ln(F_{ij}) = \beta_0 + (\gamma \cdot \ln(F_{0j})) + (\beta_1 \cdot d_{ij}) + (\beta_2 \cdot p_{ij}) + u_j + \varepsilon_{ij} \qquad (Eq. S1)$$

We can more easily compare the alternative, more flexible model 2 (Eq. S1) to model 1 (Eq. 4 in the main text) through some simple rearrangement (Eq. S2).

$$\ln\left(\frac{F_{ij}}{F_{0j}}\right) = \beta_0 + (\gamma - 1) \cdot \ln(F_{0j}) + (\beta_1 \cdot d_{ij}) + (\beta_2 \cdot p_{ij}) + u_j + \varepsilon_{ij} \qquad (Eq. S2)$$

Here we can more clearly see that the only difference between models 1 and 2 is that γ is set equal to one in the primary model. While model 1 has a lower Akaike Information Criterion (AIC) and a lower Bayesian Information Criterion (BIC) than model 2, the differences are below the threshold of what would be considered a meaningful difference. Model 1 was selected as the primary model for this manuscript due to its relative simplicity and interpretability. However, using this alternative model is recommended for future work, especially with greater sample numbers or a wider range of initial expression values ($F_0$). Model 2 was fit using the same expectation-maximization procedure, and its coefficients were assessed using the same two-sided t-test framework with conservative degrees of freedom, as described for the primary model in the main text Methods.

Methods S2: Monolayer packing density quantification of 968 bp dsDNA strands via chemical stand melting

Monolayers of dsDNA fragments were formed on microscope slides with a layer of e-beam evaporated gold, as described in the primary Methods section. Monolayer areas were circular and 6 mm in diameter. 968 bp sfGFP gene fragments with the thiol modifier attached to the 5' end closest to the promoter region (outward orientation), with the thiol modifier attached to the 5' end furthest from the promoter region (inward orientation), or without a thiol modifier were used. The incubation concentrations of all three gene fragment variants were measured using a Qubit 3.0 (Thermo Fisher) fluorometer running v1.02 of the instrument software was used with the Qubit dsDNA HS Assay Kit. All monolayer areas were backfilled with MCH. Four replicate monolayers were produced for each monolayer type.

Strands within the monolayers were dehybridized by applying 40 µl of 1 M NaOH to the monolayer areas and allowed to sit at room temperature for 10 minutes. Incubation under 1 M NaOH was selected as the chemical method most likely to result full monolayer dehybridization based on results obtained by Wang et al.[1] The resulting dehybridized ssDNA in NaOH was quantified using a Qubit ssDNA Assay Kit. Any readings that appeared below Qubit's limit of quantification, and thus returned as "out of range," was treated as a concentration of 0 ng/µl for the purpose of density calculations. It should be noted that this assay does not distinguish between ssDNA and dsDNA.[2] Total packing density for each monolayer was determined using the measured dehybridization concentrations, the NaOH incubation volume, and the known monolayer surface areas.

### Supplementary References


1. X. Wang, H. J. Lim, and A. Son, “Characterization of Denaturation and Renaturation of DNA for DNA Hybridization,” *Environmental Health and Toxicology 29* (2014): e2014007. https://doi.org/10.5620/eht.2014.29.e2014007.
2. Y. Nakayama, H. Yamaguchi, N. Einaga, and M. Esumi, “Pitfalls of DNA Quantification Using DNA-Binding Fluorescent Dyes and Suggested Solutions,” *PLOS ONE 11*, no. 3 (2016): e0150528. https://doi.org/10.1371/journal.pone.0150528.
